# Host-Specific Fluorescence Dynamics in Legume-Rhizobia Symbiosis During Nodulation

**DOI:** 10.1101/2025.10.11.681774

**Authors:** Chandan K. Gautam, Gayathri Senanayake, Amanda B. Pease, Mohamed A. Salem, Ahmed H. Rabia, Birgit M. Prüß, Barney A. Geddes

## Abstract

The legume-rhizobia symbiosis is a cornerstone of sustainable agriculture due to its ability to facilitate biological nitrogen fixation. Still, real-time visualization and quantification of this interaction remain technically challenging, especially across different host backgrounds. In this study, we systematically evaluate the efficacy of the nitrogenase system *nifH* promoter (P*nifH*) in driving expression of distinct fluorescent reporters; superfolder yellow fluorescent protein (sfYFP), superfolder cyan fluorescent protein (sfCFP), and various red fluorescent proteins (RFPs) within root nodules of determinate (*Lotus japonicus-Mesorhizobium japonicum*) and indeterminate (*Pisum sativum-Rhizobium leguminosarum*) systems. We show that P*nifH*-driven sfYFP and sfCFP yield strong, uniform, and reproducible fluorescence in nodules of both systems, facilitating reliable quantification of nodulation traits and strain occupancy. In contrast, RFPs including monomeric (mScarlet-I, mRFP1, mARs1) and multimeric (AzamiRed1.0) variants exhibited weak or inconsistent signals in pea. Notably, fluorescent labeling did not impair rhizobial competitiveness for root nodule occupancy, and P*nifH*-driven sfYFP and sfCFP reporters enabled robust multiplexed imaging in single-root and split-root assays. In the lotus, mScarlet-I worked robustly and facilitated a tripartite strain labeling system. Complementing our molecular toolkit, we established a deep learning-based analytical pipeline for high-throughput, automated quantification of nodulation traits, validated against standard ImageJ analysis. Altogether, our results identify P*nifH*-driven sfYFP and sfCFP as robust, broadly applicable reporters for legume-rhizobia symbiosis studies, while highlighting the need for optimized red fluorophores in some contexts. The integration of validated promoter-reporter constructs with state-of-the-art computational approaches provides a scalable framework for dissecting the spatial and competitive dynamics of plant-microbe mutualisms.

**IMPORTANCE:** The legume-rhizobia symbiosis is central to sustainable agriculture through its capacity for biological nitrogen fixation, yet tools for real-time, quantitative visualization of this interaction remain limited. Here, we demonstrate that the *nifH* promoter (P*nifH*) effectively drives expression of superfolder yellow (sfYFP) and cyan (sfCFP) fluorescent proteins in both determinate (*Lotus japonicus-Mesorhizobium japonicum*) and indeterminate (*Pisum sativum-Rhizobium leguminosarum*) nodules. These reporters enable robust, reproducible fluorescence without impairing rhizobial competitiveness, supporting multiplexed imaging and quantitative nodulation analyses. By contrast, red fluorescent proteins exhibited host-dependent variability, underscoring the need for improved red fluorophores. Integration of validated promoter-reporter constructs with a deep learning-based image analysis pipeline establishes a scalable framework for high-throughput assessment of nodule occupancy and symbiotic dynamics. This work provides a practical molecular and computational toolkit for dissecting plant-microbe mutualisms across diverse host systems.

## Introduction

The symbiotic interaction between legumes and rhizobia is governed by complex genetic and molecular mechanisms (1). Advances in molecular biology and genomics have significantly deepened our understanding of these symbiotic events. A pivotal breakthrough was the identification of three main groups of rhizobial genes (*nod, nif,* and *fix*), which revolutionized research on nodulation from the bacterial perspective (2–6). Further, with the advent of transcriptomics in nodulating legumes, researchers have been able to achieve an even broader view of the molecular events taking place during legume-rhizobia symbiosis (7, 8). Legumes produce root exudates which include flavonoids and iso-flavonoids (9–11). These compounds trigger the biosynthesis of rhizobial Nod factors (lipo-chitooligosaccharide) (12). These products of the *nod* genes trigger initial plant responses such as root hair curling and cortical cell division upon the contact with the rhizobia, thus initiating infection and nodule formation (7, 13). After successful infection, *nif* genes encode the nitrogenase enzyme complex, primarily dinitrogenase reductase, which is essential for converting atmospheric nitrogen (N_2_) to ammonia (NH ^+^) (14). The *fix* genes code for metabolic proteins required for nitrogenase function, which are regulated under the low-oxygen (microaerobic) conditions present within the nodule (15, 16).

Early studies faced significant gaps in knowledge regarding the spatial distribution and real-time monitoring of rhizobia within the legume host. This was primarily due to limitations in technology; traditional approaches, such as staining and light microscopy, could only provide static snapshots (17, 18). While the advent of gene-specific PCR fingerprinting, strain-specific nucleotide barcodes employing next-generation sequencing and metagenomic techniques improved resolution and enabled investigation of the effectiveness and competitiveness of multi-strain competition, their use remains constrained by complexity and cost (17, 19–23).

A breakthrough in studying real-time plant invasion dynamics by rhizobia was achieved with the development of green fluorescent protein (GFP) based reporters in *Sinorhizobium meliloti*, driven by the constitutive tryptophan promoter, *trp* (P*trp*) from *Salmonella typhimurium*, and the successful visualization of this reporter in the host *Medicago sativa* (alfalfa) (24). In parallel, utilizing the *nifH* nitrogenase promoter to drive the expression of the chromogenic beta-glucuronidase marker gene *gusA* within transposons introduced to *Rhizobium tropici* CIAT899, demonstrated a reliable reporter of nitrogen fixation in *Phaseolus vulgaris L. cv. Riz 30* (common bean) (25). This system was useful for comparing the competitiveness of labeled strains to unlabeled isogenic parents. The system was later improved to allow concurrent labeling of two different strains by the inclusion of an alternate thermostable beta-galactosidase-encoding gene, *celB*, and implemented in the *R. leguminosarum-Pisum sativum* system (26). Although these studies utilized the *nifH* promoter system (specific to nitrogen fixation), the staining procedure for chromogenic markers terminated cellular activity, preventing live imaging. In later studies, *S. typhimurium* P*trp*-driven DsRed and GFP were introduced into different strains of *S. meliloti* to study competitiveness (17, 27). However, a limitation with this reporter system was its non-specificity in reporting nodulation events. Nevertheless, this research laid the foundation for further improvement of reporter systems, which can be used explicitly to study rhizobial dynamics during nodulation within the host. Currently, *nifH* promoter (P*nifH*)-driven fluorescent reporters are the gold standard for monitoring effective nodulation and nitrogen fixation in legume-rhizobia studies (22, 25, 28, 29). The *nifH* gene encodes a key nitrogenase component and is specifically activated under the microaerobic conditions inside nodules where nitrogenase functions, making it a precise marker of active nitrogen-fixing symbiosis (25, 30, 31). Its expression is regulated by the FixJ/FixL system via FixK and NifA intermediates which are functional under microaerobic conditions, reflecting both nitrogen fixation gene activation and the low-oxygen nodule environment (3, 31–33). This specificity enables the accurate quantification of functional nodules that contribute to plant nitrogen nutrition. In contrast, *nod* gene promoters (*e.g.*, *nodABC*) are activated early in the legume-rhizobia interaction, leading to Nod factor synthesis, which triggers nodule initiation (3, 30, 34). Their activity typically decreases sharply after infection thread formation and the onset of nodule organogenesis (34). Thus, the activity of nod promoters does not reliably reflect the presence of mature, functioning nodules, and their use would result in false positives unrelated to effective nitrogen fixation. The *fix* gene promoters exhibit strong regulation by oxygen levels and can be expressed even in free-living rhizobia when exposed to low-oxygen conditions outside the plant (32, 33). Therefore, expression from these promoters is not strictly limited to symbiotic conditions, resulting in reduced specificity if used as a marker for nodulation events.

In this study, we systematically evaluated the efficacy of a P*nifH*-driven fluorescent reporter system within legume–rhizobia symbiosis by employing three spectrally distinct fluorescent proteins-superfolder yellow fluorescent protein (sfYFP), superfolder cyan fluorescent protein (sfCFP), and red fluorescent proteins (RFPs: mScarlet-I, AzamiRed1.0, mRFP1and mARs1) - each expressed under the control of the P*nifH* promoter (35–39). These constructs were tested in two model legume-rhizobia associations: *Pisum sativum* (pea)*-Rhizobium* (indeterminate nodules) and *Lotus japonicus* (lotus)*-Mesorhizobium* (determinate nodules). Our analyses revealed that sfCFP and sfYFP consistently produced robust and uniform fluorescence signals in nodules of both pea and lotus. To further benchmark reporter performance, we compared these results to those obtained using a constitutive σ-70 consensus promoter, thereby assessing the impact of promoter choice on fluorescence intensity and spatial distribution. To complement these molecular tools with scalable phenotyping, we built a deep learning (DL) workflow that converts fluorescence micrographs into quantitative readouts of symbiotic performance. By integrating spectral discrimination with high-throughput, quantitative phenotyping, our approach not only advances the toolkit for studying legume–rhizobia interactions but also opens new avenues for dissecting the spatial and competitive dynamics of symbiotic partnerships. This scalable framework has the potential to transform how we interrogate plant–microbe mutualisms, enabling deeper, system-level insights into the factors that govern symbiotic efficiency and resilience.

## Materials and Methods

### Bacterial strains and growth conditions

Plasmid constructs carrying promoter-driven fluorescent protein-encoding genes were assembled in *Escherichia coli.* Cultures of *E. coli* were maintained at 37 °C in Luria-Bertani (LB) medium (40) supplemented with appropriate antibiotics (Table 1). Construct assembly was carried out using a BsaI-based Golden Gate cloning strategy, incorporating either the P*nifH* promoter or synthetic universal promoters (P*uni*). The P*uni* included the *E. coli* constitutive σ-70 consensus J23106 (promoter) and B0032m (ribosome binding site) from the CIDAR toolkit (41), which was originally derived from the Anderson promoter library. Plasmid components, including the fluorescent protein sequences, were obtained from Addgene (https://www.Addgene.org/) or synthesized by commercial providers and subsequently assembled in-house. All constructs were sequence-verified in *E. coli* before mobilization into rhizobial recipient strains via conjugation. The rhizobial strains used in this study are *R. leguminosarum* bv. *viciae* 3841(42) (reclassified as *R. johnstonii,*(43)) and *M. japonicum* strain R7A (formerly known as *M. loti*). Additionally, two novel *Rhizobium* strains, G22 and G11, isolated from North Dakota (USA), were also used in this study. The fluorescent proteins used to label rhizobia are: sfYFP, sfCFP, mScarlet-I, mARs1, AzamiRed1.0 (Azami Red) and mRFP1 (35–39). The details of the plasmids and the sequence of fluorescent proteins are presented in Table 1 and S1, as well as in the accompanying Microbiology Resource Announcements (MRA) publications: Senanayake et al. (2025) and Pease et al. (2025). Both rhizobia strains were cultivated at 30°C in tryptone-yeast (TY) medium, a complex, non-selective growth medium suitable for a broad range of bacteria (44), supplemented with 2.5 mM CaCl_2_ and appropriate antibiotics (Senanayake et al., 2025; Pease et al., 2025; and Table 1). The resulting fluorescently labeled rhizobia were used for all plant inoculation experiments.

**Table 1:** Details of the plasmid components and constructs, and plasmid-engineered rhizobia used in this paper. Further details can be found in the Microbiology Resource Announcement (MRA) manuscripts by Senanayake et al., 2025, and Pease et al., 2025.

| Name | Description | Source | Addgene Plasmid # |
| --- | --- | --- | --- |
| C3m | superfolder green fluorescent protein (sfGFP), Golden Gate component;<br>Addgene CIDAR MoClo Vol.1<br>Extension (kit #1000000161); Amp<sup>R</sup> | Richard Murray<br>Lab: CIDAR<br>MoClo Extension<br>Unpublished | 120956 |
| C5m | monomeric red fluorescent protein; mRFP1, Golden Gate component;<br>Addgene CIDAR MoClo Vol.1<br>Extension (kit #1000000161); Amp<sup>R</sup> | Richard Murray<br>Lab: CIDAR<br>MoClo Extension<br>Unpublished | 120957 |
| C51m | superfolder yellow fluorescent protein (sfYFP), Golden Gate component;<br>Addgene CIDAR MoClo Vol.1<br>Extension (kit #1000000161); Amp<sup>R</sup> | Richard Murray<br>Lab: CIDAR<br>MoClo Extension<br>Unpublished | 120975 |
| C91m | superfolder cyan fluorescent protein (sfCFP), Golden Gate component;<br>Addgene CIDAR MoClo Vol.1<br>Extension (kit #1000000161); Amp<sup>R</sup> | Richard Murray<br>Lab: CIDAR<br>MoClo Extension<br>Unpublished | 120999 |
| C99m | mScarlet-I, Golden Gate component; | Richard Murray | 121003 |
|  | Addgene CIDAR MoClo Vol.1<br>Extension (kit #1000000161); Amp<sup>R</sup> | Lab: CIDAR<br>MoClo Extension<br>Unpublished |  |
| J23106 | J23106 Anderson library promoter,<br>Golden Gate component; Addgene<br>CIDAR MoClo Parts Kit (kit<br>#1000000059); Amp<sup>R</sup> | (41) | 65992 |
| BCD12 | BCD12 bicistronic ribosome binding<br>cite, Golden Gate component; Addgene<br>CIDAR MoClo Parts Kit (kit<br>#1000000059); Amp<sup>R</sup> | (41) | 66023 |
| pOGG043 | Synthetic *Rhizobium PnifH* Golden Gate<br>component Rlv_PsnifH; Sp<sup>R</sup> | (22) | 133123 |
| pNDGG003 | BEVA 2.0 level 1 Golden Gate cloning<br>vector with Bsa1 exchangeable lacZ,<br>RK2origin of replication, and par<br>stability, Tc<sup>R</sup> | (50) | 231316 |
| pNDGG004 | BEVA 2.0 level 1 Golden Gate cloning<br>vector with Bsa1 exchangeable lacZ,<br>pBBR1origin of replication, and par<br>stability, Tc<sup>R</sup> | (50) | 231317 |
| pNDGG037 | BEVA 2.0 T2m terminator with DF<br>extension for CIDAR MoClo GG; Sp<sup>R</sup> | (50) | 231337 |
| pNDGG076 | Synthetic *Mesorhizobium PnifH* PU/AB<br>Golden Gate component Meso_PsnifH;<br>Amp<sup>R</sup> | This work |  |
| pNDGG078 | mARs1 SC/CD Golden Gate component;<br>Amp<sup>R</sup> | This work |  |
| pNDGG079 | AzamiRed1.0 SC/CD Golden Gate<br>component; Amp<sup>R</sup> | This work |  |
| pNDMS054 | Level 1 Golden Gate Cloning with<br>pNDGG003 as backbone with parts:<br>J23106-BCD12-sfGFP-T2m; Tc<sup>R</sup> | (50) |  |
| pNDMS055 | Level 1 Golden Gate Cloning with<br>pNDGG003 as backbone with parts:<br>J23106-BCD12-sfCFP-T2m; Tc<sup>R</sup> | (50) |  |
| pNDMS056 | Level 1 Golden Gate Cloning with<br>pNDGG003 as backbone with parts:<br>J23106-BCD12-sfYFP-T2m; Tc<sup>R</sup> | (50) |  |
| pNDMS057 | Level 1 Golden Gate Cloning with<br>pNDGG003 as backbone with parts:<br>J23106-BCD12-mScarlet-I-T2m; Tc<sup>R</sup> | (50) |  |
| pNDMS156 | Level 1 Golden Gate Cloning with<br>pNDGG003 as backbone with parts:<br>Meso_PsnifH-sfGFP-T2m; Tc<sup>R</sup> | Accompanying<br>manuscript<br>Senanayake et al.<br>2025 |  |
| pNDMS157 | Level 1 Golden Gate Cloning with<br>pNDGG004 as backbone with parts:<br>Meso_PsnifH-sfGFP-T2m; Tc<sup>R</sup> | Accompanying<br>manuscript<br>Senanayake et al.<br>2025 |  |
| pNDMS237 | Level 1 Golden Gate Cloning with<br>pNDGG004 as backbone with parts:<br>Meso_PsnifH-sfYFP-T2m; Tc<sup>R</sup> | Accompanying<br>manuscript<br>Senanayake et al.<br>2025 |  |
| pNDMS238 | Level 1 Golden Gate Cloning with<br>pNDGG004 as backbone with parts:<br>Meso_PsnifH-sfCFP-T2m; Tc<sup>R</sup> | Accompanying<br>manuscript<br>Senanayake et al.<br>2025 |  |
| pNDMS239 | Level 1 Golden Gate Cloning with<br>pNDGG004 as backbone with parts:<br>Meso_PsnifH-mScarlet-I-T2m; Tc<sup>R</sup> | Accompanying<br>manuscript<br>Senanayake et al.<br>2025 |  |
| pNDMS290 | Level 1 Golden Gate Cloning with<br>pNDGG004 as backbone with parts:<br>Rlv_PsnifH-sfYFP-T2m; Tc<sup>R</sup> | Accompanying<br>manuscript Pease et<br>al. 2025 |  |
| pNDMS291 | Level 1 Golden Gate Cloning with<br>pNDGG004 as backbone with parts:<br>Rlv_PsnifH-sfCFP-T2m; Tc<sup>R</sup> | Accompanying<br>manuscript Pease et<br>al. 2025 |  |
| pNDMS292 | Level 1 Golden Gate Cloning with | Accompanying |  |
|  | pNDGG004 as backbone with parts:<br>Rlv_PsnifH-mScarlet-I-T2m; Tc<sup>R</sup> | manuscript Pease et al. 2025 |  |
| pNDMS377 | Level 1 Golden Gate Cloning with<br>pNDGG004 as backbone with parts:<br>J23106-BCD12-sfYFP-T2m; Tc<sup>R</sup> | This work |  |
| pNDMS378 | Level 1 Golden Gate Cloning with<br>pNDGG004 as backbone with parts:<br>J23106-BCD12-sfCFP-T2m; Tc<sup>R</sup> | This work |  |
| pNDMS379 | Level 1 Golden Gate Cloning with<br>pNDGG004 as backbone with parts:<br>J23106-BCD12-mScarlet-I-T2m; Tc<sup>R</sup> | This work |  |
| pNDMS344 | Level 1 Golden Gate Cloning with<br>pNDGG004 as backbone with parts:<br>Rlv_PsnifH-mRFP1-T2m; Tc<sup>R</sup> | This work |  |
| pNDMS803 | Level 1 Golden Gate Cloning with<br>pNDGG004 as backbone with parts:<br>Rlv_PsnifH-mARs1-T2m; Tc<sup>R</sup> | This work |  |
| pNDMS805 | Level 1 Golden Gate Cloning with<br>pNDGG004 as backbone with parts:<br>Rlv_PsnifH-AzamiRed1.0-T2m; Tc<sup>R</sup> | This work |  |
| pNDMS807 | Level 1 Golden Gate Cloning with<br>pNDGG004 as backbone with parts:<br>J23106-BCD12-mRFP1-T2m; Tc<sup>R</sup> | This work |  |
| pNDMS808 | Level 1 Golden Gate Cloning with<br>pNDGG004 as backbone with parts:<br>J23106-BCD12-mARsI-T2m; Tc<sup>R</sup> | This work |  |
| pNDMS809 | Level 1 Golden Gate Cloning with<br>pNDGG004 as backbone with parts:<br>J23106-BCD12-AzamiRed1.0-T2m; Tc<sup>R</sup> | This work |  |
| RmND298 | pNDMS156 conjugated into *M.*<br>*japonicum* strain R7A; Tc<sup>R</sup> | This work |  |
| RmND299 | pNDMS157 conjugated into *M.*<br>*japonicum* strain R7A; Tc<sup>R</sup> | This work |  |
| RmND533 | pNDMS237 conjugated into *M.*<br>*japonicum* strain R7A; Tc<sup>R</sup> | This work |  |
| RmND534 | pNDMS238 conjugated into *M.*<br>*japonicum* strain R7A; Tc<sup>R</sup> | This work |  |
| RmND535 | pNDMS239 conjugated into *M.*<br>*japonicum* strain R7A; Tc<sup>R</sup> | This work |  |
| RmND696 | pNDMS808 conjugated into *R.*<br>*leguminosarum* strain Rlv3841; Sm<sup>R</sup>,Tc<sup>R</sup> | This work |  |
| RmND697 | pNDMS809 conjugated into *R.*<br>*leguminosarum* strain Rlv3841; Sm<sup>R</sup>,Tc<sup>R</sup> | This work |  |
| RmND698 | pNDMS807 conjugated into *R.*<br>*leguminosarum* strain Rlv3841; Sm<sup>R</sup>,Tc<sup>R</sup> | This work |  |
| RmND700 | pNDMS803 conjugated into *R.* | This work |  |
|  | *leguminosarum* strain Rlv3841; Sm<sup>R</sup>,Tc<sup>R</sup> |  |  |
| RmND701 | pNDMS344 conjugated into *R.*<br>*leguminosarum* strain Rlv3841; Sm<sup>R</sup>,Tc<sup>R</sup> | This work |  |
| RmND702 | pNDMS805 conjugated into *R.*<br>*leguminosarum* strain Rlv3841; Sm<sup>R</sup>,Tc<sup>R</sup> | This work |  |
| RmND718 | pNDMS379 conjugated into *R.*<br>*leguminosarum* strain Rlv3841; Sm<sup>R</sup>,Tc<sup>R</sup> | This work |  |
| RmND719 | pNDMS377 conjugated into *R.*<br>*leguminosarum* strain Rlv3841; Sm<sup>R</sup>,Tc<sup>R</sup> | This work |  |
| RmND720 | pNDMS378 conjugated into *R.*<br>*leguminosarum* strain Rlv3841; Sm<sup>R</sup>,Tc<sup>R</sup> | This work |  |
| RmND623 | pNDMS290 conjugated into *R.*<br>*leguminosarum* strain Rlv3841; Sm<sup>R</sup>,Tc<sup>R</sup> | This work |  |
| RmND624 | pNDMS291 conjugated into *R.*<br>*leguminosarum* strain Rlv3841; Sm<sup>R</sup>,Tc<sup>R</sup> | This work |  |
| RmND625 | pNDMS292 conjugated into *R.*<br>*leguminosarum* strain Rlv3841; Sm<sup>R</sup>,Tc<sup>R</sup> | This work |  |
| RmND635 | pNDMS290 conjugated into *Rhizobium*<br>strain G22; Tc<sup>R</sup> | This work |  |
| RmND636 | pNDMS291 conjugated into *Rhizobium*<br>strain G22; Tc<sup>R</sup> | This work |  |
| RmND629 | pNDMS290 conjugated into *Rhizobium*<br>strain G11; Tc<sup>R</sup> | This work |  |

### Design of custom synthetic Mesorhizobium nifH promoter

For *Rhizobium*, the synthetic consensus promoter Rlv P*nifH* described by Mendoza-Suárez et al., 2020 (22) was used to drive different fluorophores (Table S1).

For *M. japonicum,* a custom synthetic *Mesorhizobium nifH* promoter was designed and implemented based on multiple sequence alignment of *nifH* upstream regions from reference strains of several symbiotic *Mesorhizobium* species. These included *M. amorpha* CCNWGS0123, *M. ciceri* bv. *biserrulae* WSM1284 and WSM1271, *M. jarvisii* ATCC33669, *M. ciceri* CC1192A, *M. muleiense* AHPC30, *M. loti* TONO, and *M. loti* MAFF303099. Consensus sequences from a region of high homology that spanned 295 bp upstream of the *nifH* coding sequence were used for the design of the synthetic sequence (Fig. S1, Table S1).

### Seed sterilization and germination

*P. sativum* CDC Striker (45) field peas and *L. japonicus cv.* Gifu seeds were used in this study. Pea seeds were surface sterilized in 50 mL Falcon tubes using sequential treatments: 95% ethanol (30s) followed by 2% sodium hypochlorite (5min), with ten sterile ddH O rinses. Sterilized seeds were germinated on 0.7% (w/v) agar plates in darkness for 48h. Lotus seeds were scarified with concentrated sulfuric acid (15 mL, 10–12 min), rinsed 10 times with sterile ddH O, then sterilized in 2.5% sodium hypochlorite (2 min) followed by ten washes with sterile ddH O. After soaking in ddH O for 4h, seeds were germinated on 0.8% (w/v) water agar plates wrapped in foil and incubated in darkness at room temperature for 48h.

### Plant growth and inoculation

The germinated seedlings of pea and lotus were cultivated in Leonard jars filled with a 1:1 (w/w) mixture of quartz sand and vermiculite. Each jar was supplemented with 250 mL of sterile nitrogen-free rooting solution (Table S2) for pea or with Jensen’s medium supplemented with 0.5 mM KNO for lotus (Table S3) (61). Plants were maintained in a controlled growth chamber under long-day conditions. For pea, conditions consisted of an 18-hour light/6-hour dark photoperiod with day/night temperatures of 21°C/17 °C. Lotus was grown under a 16 h light/8 h dark cycle at a constant temperature of 20 °C.

Three days after sowing, seedlings were inoculated with 10 mL of the respective rhizobial suspension expressing fluorescent proteins under either the P*nifH* or P*uni* promoter. Pea seedlings were inoculated with *Rhizobium* (OD = 0.06), while lotus seedlings received *M. japonicum* (OD = 0.08) inoculum.

The co-inoculation experiments were performed independently for *Rhizobium,* and *M. japonicum* on their respective hosts, pea and lotus. Equal volumes of the respective rhizobial cultures (with similar OD_600_ values) with unique fluorescent labels were mixed to yield a total volume of 10 mL, which was then applied to the seedlings in a Leonard jar. The final OD of the inoculum was same as above, *Rhizobium* (OD = 0.06), and *M. japonicum* (OD = 0.08).

### Split-root system preparation

For split-root experiments, pea seeds were surface-sterilized and germinated on sterile 0.7% agar plates, as described above. The germinated seedlings were excised at the junction of the hypocotyl and radicle using a sharp sterile blade, inside a laminar hood. The seedlings were then carefully transferred into the holes of a 1 mL pipette tip box filled with sterile nitrogen-free ½- strength Hoagland nutrient solution (46) (Table S4), ensuring that the cut region remained in contact with the nutrient solution to prevent dehydration (Fig. S2). The lid of the tip box was loosely placed on top, and the setup was maintained in a growth chamber under a photoperiod of 18h light at 21°C and 6h dark at 17°C for 4-5 days to allow the formation of adventitious roots.

After this period, all but two morphologically similar lateral roots were carefully removed using precision scissors inside the laminar hood. The remaining two roots were positioned such that each extended into a separate hole of the same tip box, allowing for individual development. The lid was kept loosely closed to avoid shoot constriction and minimize damage from condensed water. After another 12-14 days of growth, plants with clearly separated elongated-root systems were removed, and the split-roots were transplanted into separate Leonard jars (placed side by side, as shown in Fig. S2) containing a 1:1 (w/w) mixture of quartz sand and vermiculite, along with 250 mL of sterilized nitrogen-free rooting solution (Table S2, Fig. S2) (47). The plants were kept in a growth chamber for hardening for 2 days before being subjected to various treatments. The complete workflow is illustrated in Fig. S2.

### Sample harvesting and imaging

Pea roots (and split-roots) were harvested three weeks post-inoculation (*wpi*), while lotus roots were collected five *wpi*. After carefully washing the samples in water, whole plants were stored at 4°C with roots wrapped in damp paper towels until imaging. Prior to imaging, roots were trimmed to remove non-nodulated regions and arranged on a single-well cell culture plate (pea) or a microscope slide (lotus).

For pea, fluorescence microscopy was conducted using a *Biotek Cytation™ 5* (version 3.10) equipped with a 1.25X PL APO objective, while for lotus 4X PL APO was used. Image acquisition settings for Bright Field (BF) and fluorescence channels, with excitation/emission wavelengths optimized for YFP (500, 542 nm), RFP (530, 593 nm), and CFP (445, 510 nm), were tailored for each bioreporter. For P*nifH*::fluorescent protein imaging, LED intensities were set to BF 3, YFP 10, RFP 10, and CFP 10, with integration times of BF 100 ms, YFP 5 ms, RFPs ranging from 24 to 157 ms depending on the protein (mRFP1: 116 ms, AzamiRed1.0: 157 ms, mARS1: 24 ms, mScarlet-I: 24 ms), and CFP at 5 ms. Gain settings were BF 0.8, YFP 19.5, RFPs 24 (mRFP1: 24, AzamiRed1.0: 24, mARS1: 24, mScarlet-I: 24), and CFP 24.

For P*uni*::fluorescent protein imaging, LED intensities were identical (BF 3, YFP 10, RFP 10, CFP 10), while integration times were BF 100 ms, YFP 40 ms, RFPs ranging from 24 to 157 ms (mRFP1: 72 ms, AzamiRed1.0: 157 ms, mARS1: 30 ms, mScarlet-I: 24 ms), and CFP 12 ms.

Gain settings were set to BF 0, YFP 20, RFPs ranging between 24 and 30 (mRFP1: 27, AzamiRed1.0: 24, mARS1: 30, mScarlet-I: 24), and CFP 20. Focus was set either using autofocus or fixed at a height of 1000 μm with a bottom elevation of 0.2 μm. Since P*uni*-driven fluorophores don’t require a nodule-like environment to be functional, we used these settings to visualize them on a TY agar plate (Fig. S3A, B).

For lotus, image capture settings were: i) P*nifH*::fluorescent protein- (LED: BF 9, YFP 10, RFP 2, and CFP 3; integration time: BF 5, YFP 5, RFP 5, CFP 5; gain: BF 0.8, YFP 3.17, RFP 1, CFP 1) ; ii) P*uni*::fluorescent protein- (LED: BF 9, YFP 10, RFP 10, and CFP 10; integration time: BF 5, YFP 30, RFP 45, CFP 17; gain: BF 0.8, YFP 20, RFP 24, CFP 24).

### Nodule counting and quantification of nodulation parameters

The images obtained from the *Cytation^TM^ 5* were subjected to visual quantification of fluorescent and non-fluorescent nodules, as well as ImageJ-based quantification.

Fluorescent nodule number and intensity quantification using ImageJ:

- Auto-nodule detection and counting: File > Open > Load Pic (you can drag to the ImageJ floating bar) Image > Adjust > Color balance > Blue/Yellow (do it one by one for each color) > (set minimum, maximum, brightness levels based on the image needs)

► Click ‘apply’

i. Image > Adjust > Color threshold > Blue/Yellow > (set Hue, Saturation, and Brightness levels based on the image) Analyze > Analyze particles > Click > Size (pixel, to define size range) > Click OK to run the analysis **Note:** Some of the clumped nodules may be missed or counted as a single unit. Therefore, perform manual counting by following the instructions below.
- Assisted-nodule number and fluorescent intensity counting: File > Open > Load Pic (you can drag to the ImageJ floating bar) Plugin > Analyze > Cell Counter Right-click on the section of nodules where fluorescence is maximum. Use Ctrl + mouse cursor to zoom in and out. alt + mouse right click= select for deletion alt + M= result outcomes

### Data visualization and statistical analyses

All statistical analyses of nodule count data were performed using the built-in analysis tools of GraphPad Prism (version 10.5.0). Depending on the experimental design, either one-way or two-way ANOVA was applied. Post-hoc comparisons were carried out using Tukey’s Honestly Significant Difference (TukeyHSD) test, and results were summarized using a compact letter display to represent all pairwise comparisons. Data visualization, including all plots and graphs, was also generated using GraphPad Prism (version 10.5.0).

### Deep learning-based quantification of nodulation traits

To enable automated, high-throughput detection and spatial quantification of nodulation events, we developed a deep learning (DL) pipeline utilizing instance segmentation and post-inference morphometric analysis (48). This pipeline was designed to detect fluorescent nodules, classify them by reporter color, quantify their area and brightness, and evaluate their spatial relationship to the main root.

Initial training images were manually annotated using Roboflow to label fluorescent nodules according to fluorophore class: *Cyan* (sfCFP), *Yellow* (sfYFP), and *Red* (mScarlet-I and variants). A total of 16 representative images were collected, 8 of which were manually segmented and augmented to generate a 15-image training set. The remaining images were split into 4 for validation and 4 for testing to assess model performance. The YOLOv8s-seg model was then fine-tuned using these annotations (49). The model was trained for 100 epochs with the AdamW optimizer with a learning rate of 0.001429 and momentum of 0.9 to optimize detection accuracy across both model systems.

### Remote inference and postprocessing

For image analysis, we employed the Roboflow Inference SDK, querying the trained model remotely using the InferenceHTTPClient. Images were first converted to high-quality JPEGs to ensure compatibility and minimize artifacts. The inference returned class labels, bounding boxes, and confidence scores for each predicted nodule. Predictions were filtered using a confidence threshold of 0.3 to exclude spurious detections. The API endpoint used was detect.roboflow. For each detected nodule, we quantified both its projected area and its mean fluorescence brightness.

### Brightness and area calculation

To assess fluorescence intensity, we cropped each region of interest in the original RGB image according to those bounding-box coordinates. Then we computed the per-channel mean pixel values using the Python Imaging Library (PIL). Because each reporter has a distinct emission profile, we tailored our brightness calculation accordingly: nodules expressing sfCFP (the “Blue/ Cyan” class) were characterized using only the blue-channel mean, while those expressing sfYFP or mScarlet-I (the “Yellow” and “Red” classes) were quantified by averaging their red and yellow channel means to capture their mixed-spectrum signal. By combining these morphometric and photometric measurements, we derived robust metrics of nodule size and relative reporter expression intensity, enabling direct comparisons of promoter activity and colonization efficiency across treatments. The projected area, derived directly from the model’s segmentation output, was then converted to square millimeters (mm²).

### Batch analysis and aggregation

The analysis pipeline was designed to batch-process all .png images located within the working directory, allowing for efficient evaluation of large datasets. For each image, the model predictions were parsed to compute key statistics for each nodule class, including the total number of nodules, the average and total projected area (in pixels squared), and the average and total brightness, normalized on a 0–100 scale. These metrics were then aggregated into a structured summary table using the pandas library, facilitating downstream statistical analysis and visualization. The resulting DataFrame formed the basis for the violin plots and bar charts presented in Fig. 4B, which illustrate trends in nodule size distributions, relative class frequencies, and variation in fluorescence intensity across experimental conditions.

### Root detection and distance measurement

Although the segmentation model reliably identified nodules within each image, extracting the primary root structure required classical image processing techniques. To achieve this, the images were first thresholded and subjected to morphological filtering to produce a clean binary mask of the root. The distance between each detected nodule and the primary root was then calculated using a path-based approach. This method involved three key steps: first, nodules were computationally excluded from the binary mask to isolate the root network; second, a spatial filtering step retained only those pixels located within a defined proximity to the root structure, effectively removing noise and unrelated features; and third, the shortest navigable path between each nodule and the root was computed based on the retained pixel network. This approach enabled accurate modeling of spatial relationships within the root system while avoiding the limitations of Euclidean (straight-line) distance measurements. The resulting spatial data revealed biologically meaningful patterns, including a tendency for sfCFP-tagged strains to colonize away from the main root, especially in co-inoculation experiments.

This DL-enabled workflow was essential for analyzing complex datasets, particularly in split-root co-inoculation experiments. By leveraging distinct fluorescent tags and automated classification, we quantified nodule distribution and reporter activity with high spatial and spectral resolution. The ability to batch-process images and extract both biological and signal-level metrics significantly reduced manual effort and inter-observer variability.

## RESULTS

### PnifH effectively drives reliable expression of three unique fluorophores in lotus nodules but only two in pea

To develop a multiple-output reporter system that specifically marks nodule formation, accurately reflects nitrogen fixation activity, and remains unperturbed by the imposed fitness costs prior to host infection, we assembled a suite of fluorescent reporters driven by the P*nifH* promoter. This design, similar to previously described systems (22), enables reporter activity to correlate with both bacteroid abundance and the resulting nodule size. For *Rhizobium,* a synthetic consensus *Rhizobium* P*nifH* promoter was adopted from Mendoza-Suárez et al. 2020. For *M. japonicum* we designed a custom synthetic consensus *Mesorhizobium nifH* promoter based on multiple sequence alignment of *nifH* upstream regions from reference strains from several symbiotic *Mesorhizobium* species (Fig. S1 Table S1). The promoters and fluorescent modules were combined in a medium-copy-number pBBR1 backbone that included the *par* locus, enabling stable environmental gene expression (50). These bioreporters expressed sfYFP, sfCFP, and mScarlet-I under the control of the P*nifH* promoter and were introduced to wild type *R. johnsonii* Rlv3841 and *M. japonicum* R7A strains through conjugation. These engineered strains were used to inoculate pea and lotus seedlings. The roots of the nodulated plants were removed from pots and imaged for fluorescence (Fig. 1).

**Fig. 1.**
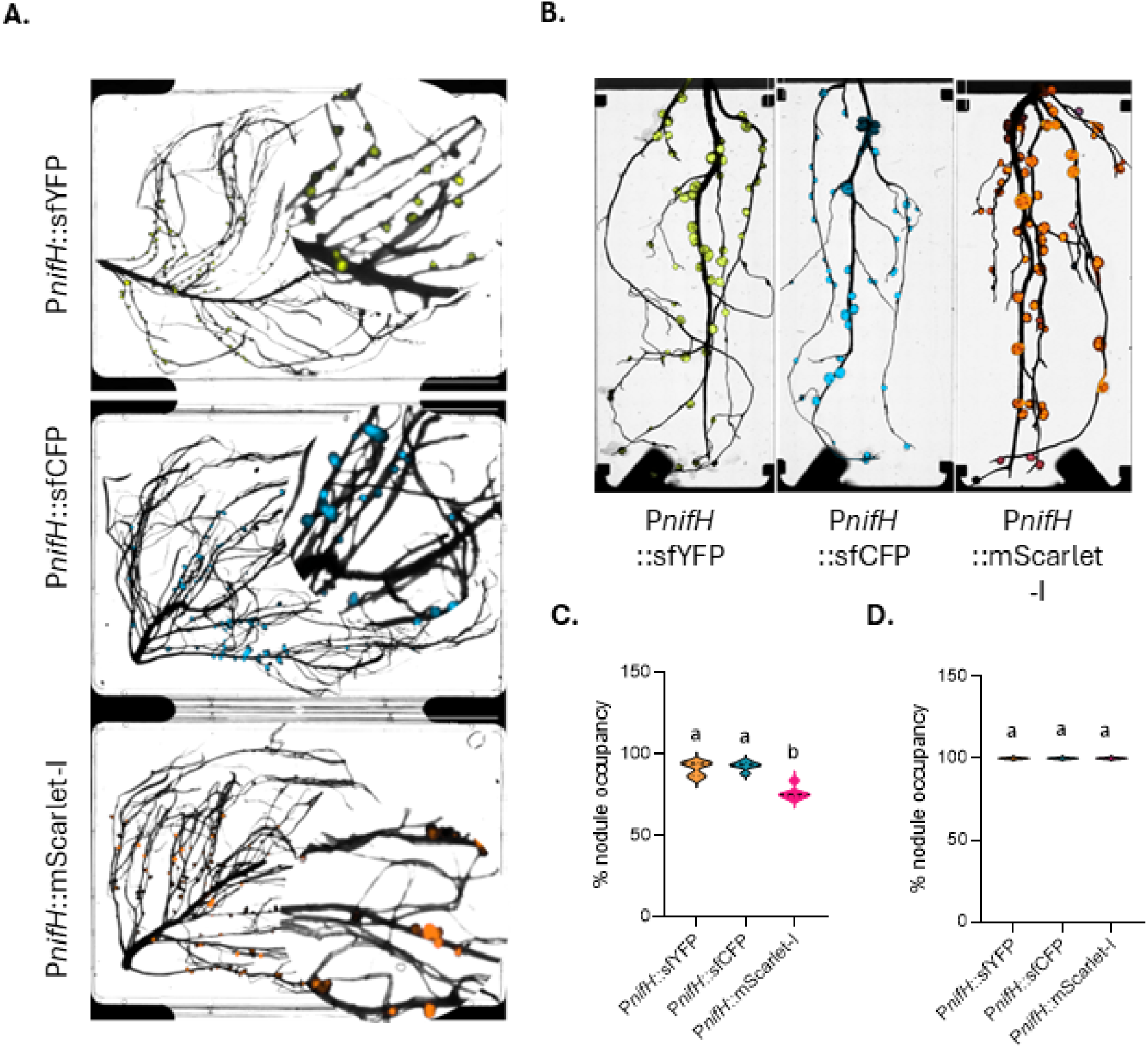
P*nifH*-driven a-pression of fluorescent proteins in *P. sativum* and *L. japonicus* nodules. (A) *Cytation^TM^5* imaging of root nodules from *P sativum* inoculated with *R. leguminosarum* strains expressing sfYFP, sfCFP, or mScarlet-1 under the control of *thePnijH* promoter. Representative images are shown from five independent plants. (B) *Cytation™ 5* imaging of root nodules from *L.japonicus* inoculated with *M japonicum* strains expressing sfYFP, sfCFP, or mScarlet-1 under the control of *thePnijH* promoter. Representative images are shown from five independent plants. (C) Percentage of fluorescently occupied nodules from *P sativum* samples shown in Fig IA. Data represent five independent biological replicates. (D) Percentage of fluorescently occupied nodules from *L.japonicus* samples shown in Fig. 1B. Data represent six to nine independent biological replicates. Distinct letters above bars indicate statistically significant differences (P < 0.05) as determined by one-way A NOVA followed by Tukey’s HSD test.

Fluorescence imaging revealed strong reporter expression in most nodules in both pea and lotus, confirming successful reporter system functionality (Fig. 1A, B). In pea, however, the proportion of fluorescent nodules was lower across all three reporter variants (sfYFP, sfCFP, and mScarlet-I) compared to lotus (Fig. 1C, D). This reduction was most pronounced for the mScarlet-I reporter, where approximately 20% of nodules in pea exhibited weak or no detectable signal (Fig. 1A, C). Adjusting imaging parameters, including increasing gain, enhanced the fluorescent signal intensity but also resulted in elevated background noise and the appearance of pseudo-signals from root tissues. Interestingly, in lotus inoculated with mScarlet-I labeled rhizobia, nearly all nodules exhibited fluorescence. Intensity and abundance of the red fluorescence were comparable to those observed with sfCFP- or sfYFP (Fig. 1B, D). Quantification of fluorescent nodules indicated a nearly consistent pattern of P*nifH*-driven expression of sfYFP and sfCFP across biological replicates in both the plant systems (Fig. 1C, D).

### Evaluation of inconsistent red fluorescent protein expression in pea nodules

In fluorescence labeling within living systems, monomeric fluorescent proteins are generally preferred due to their stability and consistent subcellular localization (39). In lotus, the monomeric RFP mScarlet-I (35), driven by synthetic P*nifH* sequences (Fig. S1B; Table S1), performed effectively (Fig. 1B, D). However, the same fluorophore showed poor performance in pea (Fig. 1A, C). We investigated whether other monomeric RFPs could enhance performance in pea. We tested monomeric RFPs such as mRFP1 and mARs1, along with a multimeric RFP, AzamiRed1.0, driven by P*nifH* (36–39). Concurrently, to evaluate if the unreliable expression of RFPs was promoter-dependent, we compared the fluorescence of these RFPs driven by a constitutive promoter (P*uni*) (41). We used a constitutive promoter to address the possibility that nodulation or bacteroid-specific physiological constraints, such as microaerobic conditions, might limit reporter performance under the P*nifH*.

Across both the P*nifH* and P*uni* promoters, all RFP variants exhibited inconsistent and generally weak fluorescence (Fig. 2A, B; S4; S5A, B). Quantification of fluorescent nodules indicated that the use of P*uni* did not change the proportion of fluorescent nodules for mARs1 and mScarlet-I but increased significantly for Azami Red and mRFP1, compared to their respective P*nifH*-driven fluorophores (Fig. 2B); nevertheless, the overall signal intensity remained low (Fig. 2A). We also evaluated the P*uni*-based fluorescence in lotus; however, in this system, it resulted in extremely low signal intensities, requiring high-gain imaging, which in turn elevated background autofluorescence in the roots (Fig. S5C).

**Fig. 2.**
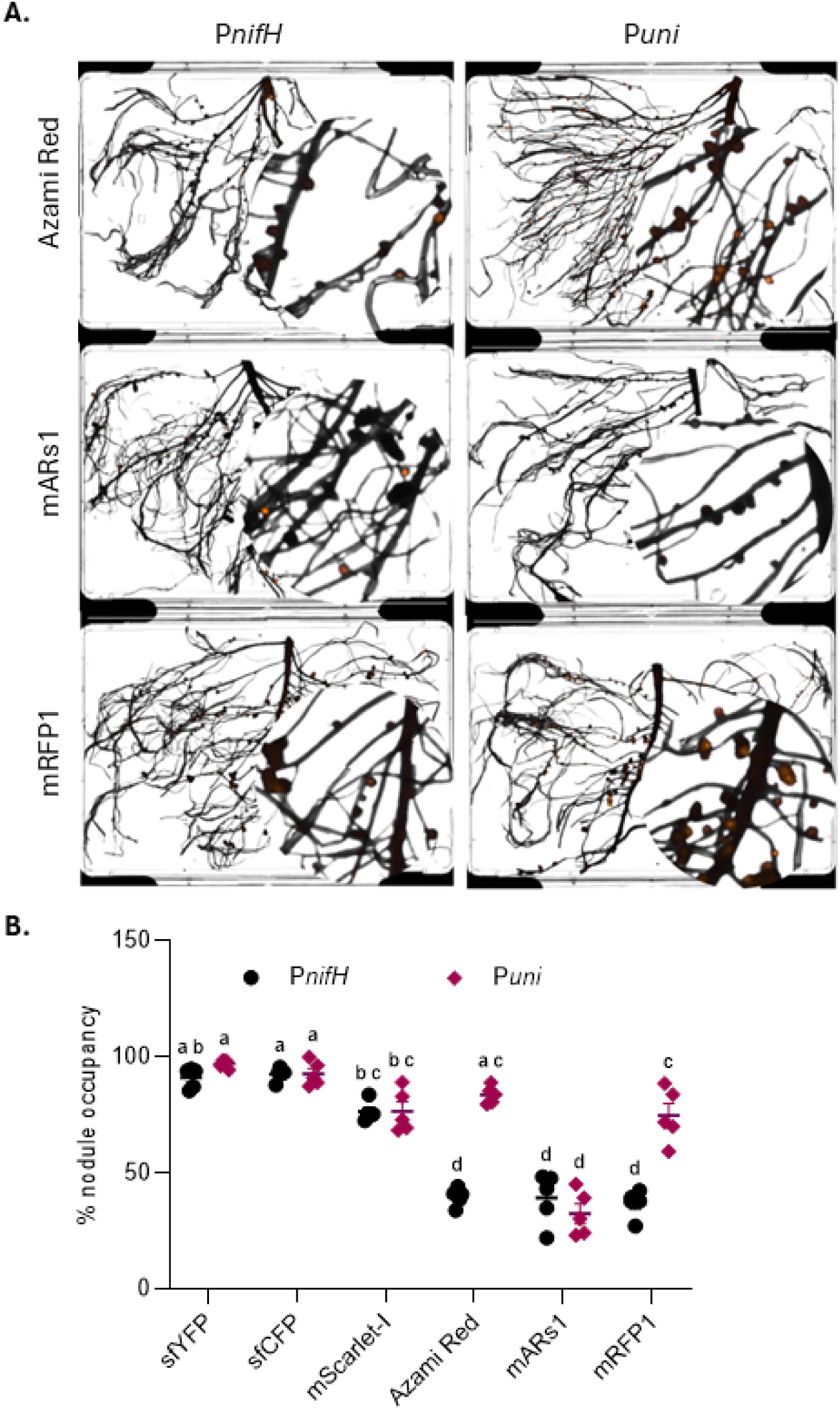
Expression of red fluorescent proteins (RFPs) in *P. sativum* nodules under P*nifH* and *Puni* promoters, and % nodule occupancy. (A) *Cytation^TM^ 5* imaging of nodules expressingAzami Red, mARsl, ormRFPl driven by either the *PnijH* or *Puni* promoter. Representative images are shown from five independent plants. (B) Percentage nodule occupancy of ***R.*** *leguminosarum* (with *PnijH* and *Puni* driven fluorescent labels) in *P sativum.* Data represent five independent biological replicates. Distinct letters above bars indicate statistically significant differences (P < 0.05) as determined by two-way ANOVAfollowed by Tukey’s HSD test.

Altogether, these data indicate that in our system, P*nifH*-driven red (mScarlet-I), cyan (sfCFP), and yellow fluorophores (sfYFP) function effectively as nodule bioreporters in the lotus-*Mesorhizobium* system, while sfYFP and sfCFP proved more reliable than mScarlet-I in the pea*-Rhizobium* system.

### Fluorescent labels do not affect rhizobia effectiveness or competitiveness when co-inoculated

A key functionality of symbiosis bioreporters is to facilitate strain identification and visualization of competitiveness for nodule formation during multi-strain co-inoculation. To determine whether P*nifH*-driven fluorescent reporters influence nodule occupancy of uniquely labeled rhizobia, we compared nodule formation by the *M. japonicum* wild type strain R7A labeled with unique fluorescent proteins, sfCFP, sfYFP, or mScarlet-I inoculated in equal ratios. When pairs of differently labeled strains were introduced together (sfCFP/sfYFP, sfCFP/mScarlet-I, or sfYFP/mScarlet-I), the percentage of nodule occupancy remained similar between the fluorescent labels (Fig. 3A, B). Even in triple inoculations with all three distinct fluorescent tags (sfCFP + sfYFP + mScarlet-I), no single reporter-containing wild-type out-competed the others, and quantification of fluorescent signal confirmed balanced representation (Fig. 3E, F). Nodules with mixed infections were readily observed at ∼10% in all combinations (Fig. S6A), in line with previous estimates (51, 52). Altogether, these findings suggest that reporter identity does not influence competitiveness in this system and that up to three distinct fluorescently labeled strains can be used simultaneously to assess nodule occupancy by *Mesorhizobia* in lotus.

**Fig. 3.**
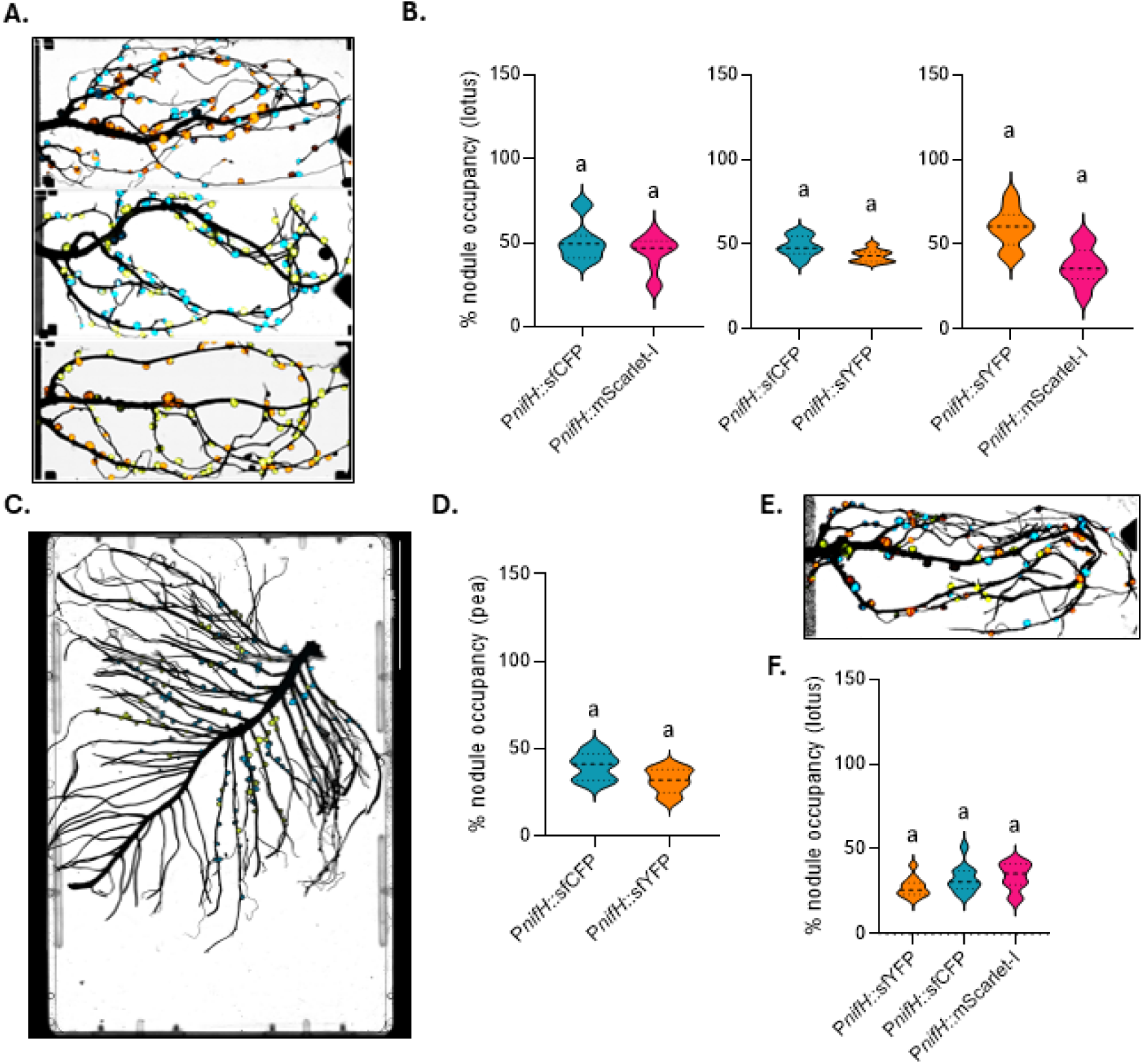
Fluorescent reporter expression in co-inoculation experiments. (A) *Cytation^TM^ 5* imaging of *L. japonicus* roots co-inoculated with equal proportions of two uniquely *labeledMjaponicum* strains (sfCFP + mScarlet-1, or sfYFP + sfCFP, or sfYFP + mScarlet-1), each expressing a *<u>PnijH</u>*-driven fluorescent protein. (B) Percentage of fluorescently occupied nodules corresponding to the samples shown in Fig. 3A. (C) *Cytation^TM^ 5* imaging of *P. sativum* co-inoculated with *R. leg11minosar11m* strains carrying *PnijH­*driven sfCFP, and sfYFP in a 1:1 ratio. (D) Percentage of fluorescently occupied nodules corresponding to the samples shown in Fig. 3C. (E) *Cytation™5* imaging of *L. japonicus* roots co-inoculated with three *M. japonicum* strains (sfCFP + sfYFP + mScarlet-1) in equal proportions, each expressing a P*nijH*-driven fluorescent protein. (F) Percentage of fluorescently occupied nodules corresponding to the samples shown in Fig. 3E. Data represent six to nine independent biological replicates for Fig. B, F, and five for Fig. D. Distinct letters above bars denote statistically significant differences (P < 0.05) based on one-way ANOVA followed by Tukey’s HSD test.

**Fig. 4.**
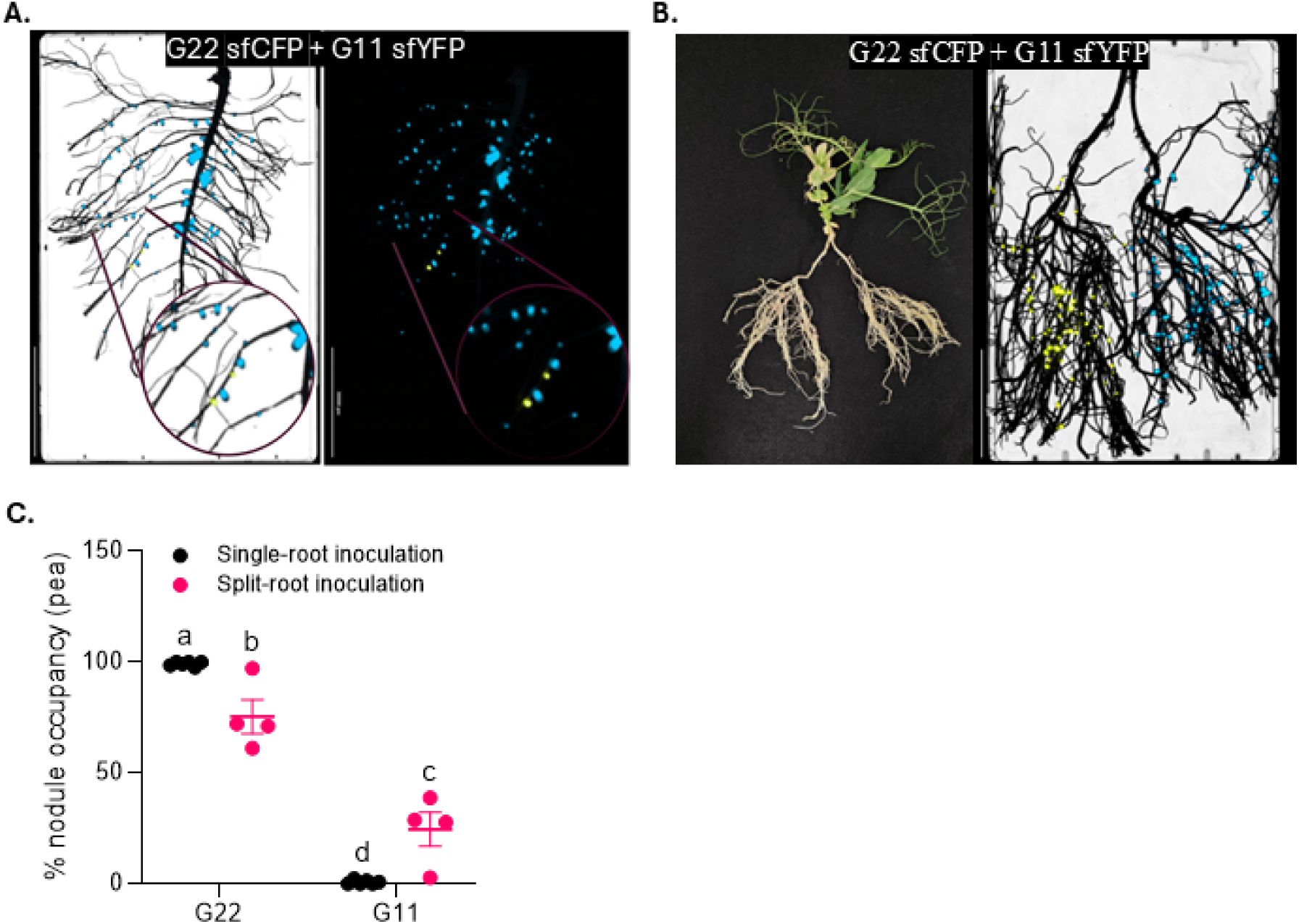
Applications of P*nifH*::fluorescent reporters in *P.sati,mm-Rhizobium* symbiosis. (A) *Cytation^TM^5* imaging of pea roots co-inoculated with equal proportions of two *Rhi=obium* strains: G22 (*PnijH*::sfCFP) and Gll (*PnijH*::sfYFP), monitored within a single-root system. (B) *Cytation™ 5* imaging of split-root plants inoculated separately with *Rhi=obium* strain G22 *(PnijH:*:sfCFP) on one root half and strain G11 *(PnijH:*:sfYFP) on the other. (C) % nodule occupancy across single-root (Fig. 4A) and split-root (Fig. 4B) systems. Data represent five independent biological replicates. Distinct letters above the plots indicate statistically significant differences (P < 0.05) as determined by two-way ANOVAfollowed by Tukey’s HSD test.

We did similar experiments with pea*-R. leguminosarum* using the wild-type strain Rlv3841 bearing bioreporters. Because P*nifH*::mScarlet-I yielded weaker, inconsistent signals in pea, we restricted analysis in this system to sfCFP and sfYFP. Similar to independent inoculations (Fig. 1A, C), there was not a significant difference in nodule occupancy between the two fluorescent labels (Fig. 3C, D). However, mixed-infection nodules were not observed (Fig. S6B) as readily as in *L. japonicus*.

### Applications of PnifH driven fluorescent reporters in pea-rhizobium symbiosis

A key utility of strain-specific bioreporters is to serve as neutral markers suitable for comparative and ecological studies (Wendlandt et al., 2025; Westhoek et al., 2017; Montoya et al., 2023). Therefore, we wished to evaluate the efficacy of the reporters described herein in these contexts. To do so, we extended this approach to assess competitiveness among naturally occurring *Rhizobium* by differentially labeling two North Dakota (USA) isolates: G22 with P*nifH*::sfCFP and G11 with P*nifH*::sfYFP. When co-inoculated onto pea roots in single-root set-up in equal proportions, the two strains were readily distinguished based on their fluorescence (Fig. 4A). Consistently, strain G22 displayed a strong competitive advantage, occupying the majority of nodules (Fig. 4A).

To examine strain behavior in the absence of direct competition on the same root system, a split-root assay (Fig. S3, 4B) was performed. Each half root was inoculated separately with either the sfCFP- or sfYFP-labeled strain. Both reporters were easily visualized, and no cross-colonization between root halves was detected (Fig. 4B). Notably, in the split-root setup, the nodule occupancy of G22 exhibited a significant reduction, whereas that of G11 showed a significant increase relative to their respective values observed in the single-root setup (Fig. 4C). Together, these experiments demonstrate that P*nifH*::fluorescent reporters provide a robust tool to monitor strain competitiveness, nodule occupancy, and spatial distribution in both single-root and split-root experimental systems.

### Deep learning-based quantification of strain effectiveness and competitiveness in split-root assays

A central aspect of characterizing legume-rhizobium interactions is the quantification of nodulation traits, including nodule number, size, area, and spatial distribution of nitrogen-fixing nodules. To further benchmark its competitive performance, we compared strain G22, labeled with P*nifH*::sfYFP, against *R. johnsonii* bv. *viciae* 3841, labeled with P*nifH*::sfCFP. Co-inoculation in a pea split-root setup (Fig. S3) enabled clear visualization of both fluorescent markers with no evidence of cross-contamination (Fig. 5A). This system thus provided a reliable platform for controlled comparisons of strain competitiveness. Across these assays, G22 again demonstrated a clear competitive advantage, occupying a majority of nodules over Rlv3841 (Fig. 5A). To quantitatively assess competitiveness and nodule parameters, we employed a DL-based image analysis pipeline to extract nodulation parameters, including nodule number, fluorescence intensity, and average size. The outputs of the deep learning tool revealed consistent trends, as summarized in Fig. 5B, demonstrating a vastly superior nodule formation ability of G22 relative to the reference wild-type strain Rlv3841, with a higher average nodule size. Meanwhile, the average fluorescence levels of nodules were similar between those formed by the two strains.

**Fig. 5.**
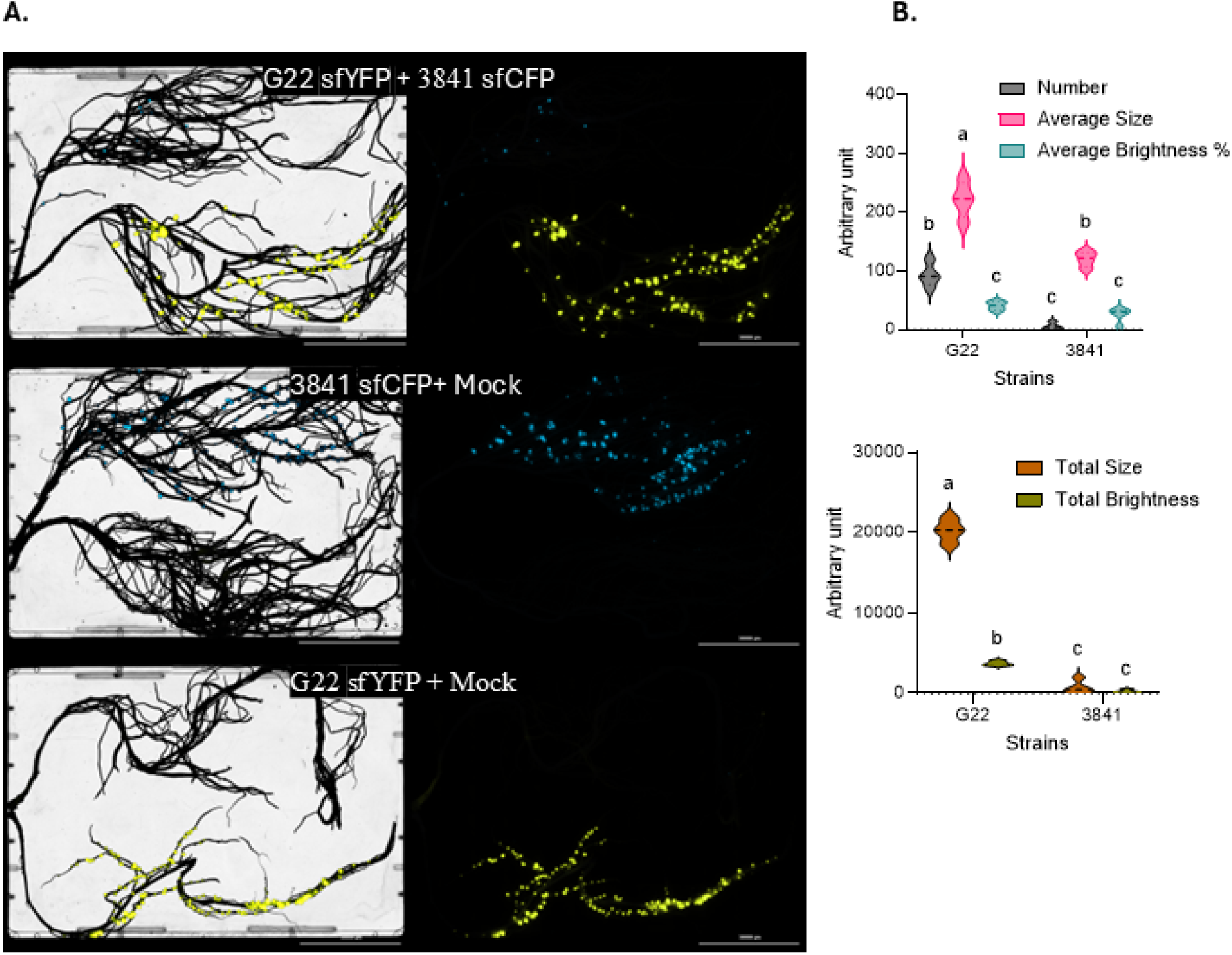
Deep-learning based quantification of nodulation traits in *P.sativum-Rhizobium*symbiosis. (A) *Cytation^TM^ 5* imaging of split-root plants inoculated with *Rhi:obium* strains G22 (P*nijH*::sfYFP) and R/v384 l *(PnifH:*:sfCFP). (B) Deep learning-based quantification of nodulation traits from split-root experiments shown in Fig. 5A. Data represent four independent biological replicates. Distinct letters above bars indicate statistically significant differences (P < 0.05) based on two-way A NOVA followed by Tukey’s HSD test.

To validate these automated results, we performed parallel quantification using conventional ImageJ-based analysis. The two approaches yielded highly concordant results, underscoring the robustness and accuracy of the DL method for nodulation trait analysis (Fig. S7). Together, these findings establish DL-driven quantification as a reliable and scalable approach for monitoring nodulation outcomes in split-root experiments, while also highlighting the power of P*nifH*::fluorescent reporters for tracing strain performance in pea*-Rhizobium* symbioses.

## Discussion

This study established newly optimized bioreporter systems for investigating legume-rhizobium symbiosis by integrating promoter-driven fluorescent reporters with automated quantification of nodulation traits. Our systematic evaluation of various P*nifH*- and P*uni*-controlled expression of fluorescent proteins across two phylogenetically distinct symbioses provided significant insights into the utility of these reporter systems for studying symbiotic nitrogen fixation. The utilization of the *nifH* promoter to control reporter expression ensures effects of free-living reporter expression do not convolute measurements of competition for nodule occupancy. Moreover, it also allows the fluorescent protein signal to act as a bioreporter of nitrogenase activity (Mendoza-Suarez et al. 2020), the utility of which is enhanced by the concurrent measurement of bioreporter activity in multiple strains on the same root. Additionally, the introduction of a high-throughput segmentation pipeline enables robust, quantitative, and non-destructive visualization of nodulation parameters, opening new possibilities for efficient and comprehensive analysis.

The P*nifH* promoter-based reporter system has become a standard tool in the study of legume-rhizobium symbiosis, reliably serving as a proxy for nitrogenase activity and as an indicator of rhizobial nodule colonization and formation of functional bacteroids (Mendoza-Suárez et al. 2020; Cebolla et al. 1994; Peralta et al. 2004). Our results indicate that P*nifH*-driven expression of sfYFP and sfCFP yields robust and reproducible fluorescence in both determinate (lotus) and indeterminate (pea) nodules, facilitating precise visualization of strain distribution, nodule occupancy, and regions of active nitrogenase activity (Fig. 1A-D; Fig. 3A-F; Fig. 4A, B). Since the P*nifH*s used in this study are derived from consensus sequences from a set of rhizobial strains, these promoter-reporters can be used in a wide range of *Mesorhizobium* and *Rhizobium* strains. Our finding reinforces the utility of P*nifH*-based fluorescent bioreporters as valuable tools for evaluating the effectiveness and competitiveness of rhizobial symbionts within diverse legume hosts.

However, not all fluorophores performed equally well. mScarlet-I exhibited robust performance in lotus (Fig. 1B, D). mScarlet-I showed inconsistent expression in pea nodules, with approximately 20% of nodules lacking an easily detectable signal (Fig. 1A, C). This was despite mScarlet-I being a monomeric protein (Shaner et al., 2004; Heppert et al., 2016), which is known to be highly stable in other biological systems. Efforts to improve mScarlet-I readability in pea nodules by adjusting imaging conditions often increased background autofluorescence, further limiting the interpretability of the results. Similar inconsistencies were observed with other RFPs, both monomeric (mRFP1, mARs1) and multimeric (Azami Red), none of which provided reliable detection in pea (Fig. 2A). These findings highlight a critical limitation in using RFPs, indicating that host-specific factors, such as the intracellular environment and nodule physiology, significantly influence reporter performance and stability. Although monomeric fluorescent proteins are generally preferred for their reliable in vivo expression in biological systems (Shaner et al. 2004; Heppert et al. 2016), the present results demonstrate that their effectiveness can be species-dependent, particularly for RFPs in pea.

To further examine reporter versatility, we tested the P*uni* promoter-based reporter system in both the legume systems, a constitutive σ-70 consensus promoter element designed for broad expression regardless of nodulation status (50, 53, 54). While P*uni*-driven fluorophores enabled analysis independent of microenvironmental cues specific to nodules, they brought limited benefits. In pea, P*uni* drove sfYFP, sfCFP, and mScarlet-I at levels similar to P*nifH*, but did not improve the reliability of RFP detection, as signals were weak and inconsistent (Fig. 2A, B; Fig. S4). Although more nodules were detected with Azami Red and monomeric mRFP1, their fluorescence was still low and variable (Fig. 2A, B). Expression from the P*uni* promoter was very weak for all fluorophores in lotus, pointing towards a host-dependent variability in expression of constitutive promoters (Fig. S5C). In indeterminant nodules of pea, where the P*uni* promoter was functional, a gradual transition between the expression of the housekeeping σ-70 *rpoD* and the σ factor associated with expressing nitrogen-fixation genes, σ-54 *rpoN,* can be observed between zones of the nodule (Roux et al. 2014). The weak expression from the P*uni* σ-70 promoter in lotus nodules may reflect a more uniform downregulation of σ-70-based gene expression within inhabitants of determinant nodules. This may have implications for promoter design for use within rhizobium bacteroids. However, it is also essential to consider that the strength of the Anderson library-derived constitutive P*uni* promoter can be further enhanced through selection of stronger variants, which may impact the efficacy of labeling in lotus nodules (Iverson et al. 2016; Bhat et al. 2024; Liow et al. 2019). Overall, these patterns underscore that the biochemical properties of both the chosen fluorescent protein and the host cellular milieu are as critical as the promoter in achieving reliable and interpretable reporter output.

Despite forming functional nodules, host-specific factors such as oxygen availability, protein folding, and chromophore maturation variables may affect fluorescent protein readouts. For instance, it is known that the maturation of GFP and RFP derivatives is oxygen-dependent (Bindels et al., 2016; Pavlou et al., 2022; Tsien, 1998). Consequently, the variability of oxygen within determinate and indeterminate nodule types may contribute to weak or inconsistent fluorescence. Determinate nodules, like those in lotus, possess an oxygen diffusion barrier at the apex, which maintains uniformly low oxygen levels throughout the nodule (Venado et al. 2022). By contrast, indeterminate nodules lack this apical barrier, resulting in a longitudinal gradient of oxygen concentration (Denison and Okano, 2003; Dixon and Kahn, 2004). We speculate that the host-specific challenges we observed with RFPs may result from differences in the oxygen levels in the pea and lotus nodules we tested in this study.

We observed other host-specific differences, including a rarer occurrence of mixed nodules in pea relative to lotus, as well as a greater proportion of “dark” nodules (Supplemental Figure S6A, B). We expected to observe a rare but significant population of mixed nodules in both systems, based on previous studies in pea (Mendoza-Suárez et al. 2020; Wendlandt et al. 2025), and lotus (Zgadzaj et al. 2015). It is possible that the P*nifH* fluorescence signal is ineffective at quantifying mixed populations in indeterminant nodules due to a biased expression in the nitrogen-fixing zone of the nodule, with the mixed strains variably occupying distinct nodule zones (Wendlandt et al. 2025). In pea, approximately 25% of nodules also did not display any fluorescent signals despite successful nodulation (Fig. S6B). The underlying cause of this observation remains unclear, but it may be associated with rhizobia exhibiting weak nitrogenase activity, potentially influenced by the anatomical and developmental characteristics of indeterminate nodules (Kereszt et al. 2011). In these nodules, the nitrogen-fixing zone is located at a distance from the meristem, resulting in a delay in the maturation of newly formed cells and a lag before rhizobia can initiate nitrogen fixation (Chaulagain and Frugoli 2021; Ferguson et al. 2010). In contrast, smaller determinate nodules develop more synchronously, leading to a more uniform and consistent rate of nitrogen fixation (Chaulagain and Frugoli 2021; Ferguson et al. 2010). This difference in developmental progression also results in uniform senescence of determinate nodules, whereas indeterminate nodules exhibit gradual senescence (Berrabah et al. 2024). Overall, these host-dependent effects reinforce the need to empirically validate fluorescent systems in each legume species, rather than assuming cross-compatibility.

Overall, our data indicates that P*nifH-*driven fluorescent proteins offer significant opportunity for studying rhizobium-legume symbiosis, with particularly promising performance within lotus nodules formed by *Mesorhizobium.* Despite the host-specific challenges we discuss above within the pea-*Rhizobium* system, we still demonstrated substantial utility of P*nifH-*sfYFP and P*nifH*:sfCFP reporters. We exemplified this utility by monitoring the fluorescence of nodules and nodulation characteristics of two natural Rhizobium strains in both single-inoculation (Fig. 4A) and split-root configurations in pea (Fig. 4B, 5), where we observed different patterns of competitive interaction (Fig. 4C). The quantification of these differences with the same set of rhizobia facilitate investigations that separate local microbe-microbe competitive interactions from plant partner choice influence on rhizobial competitiveness (55). In single-root inoculation experiments, strain G22 was highly competitive, occupying the majority of nodules (Fig. 4A, C). In contrast, under split-root conditions, where G22 and G11 were not in direct contact but exposed to shared systemic signals, nodule occupancy for G11 was significantly higher compared to its single-root inoculation condition (Fig. 4B, C). These findings underscore the significance of experimental context in assessing competitiveness and suggest that split-root assays should be incorporated as a complementary approach for a robust evaluation of rhizobial competition under diverse conditions.

Recently, the adoption of machine learning and deep learning approaches in plant biology research has accelerated markedly, particularly for quantitative trait analysis and automated phenotyping. Nonetheless, reports focusing specifically on the quantification of nodulation traits in legumes using these computational tools remain scarce. The majority of research in this domain focuses on high-throughput phenotyping platforms that characterize nodulation using image-based methods, typically applied to nodules naturally infected by rhizobia, and does not employ fluorescent bioreporters or tagged strains (56–60). To the best of current knowledge, this study represents the first report on the application of DL for analyzing fluorescent imaging data directly linked to the nitrogen fixation status of legume nodules in real time. Automated, DL-based quantification of nodule traits was integrated into split-root competition assays, enabling high-throughput and reproducible analysis that closely paralleled conventional ImageJ-based measurements, thereby validating the utility of these automated pipelines for phenotyping nodulation and symbiotic performance (Fig. 5A, B; S7).

Although P*nifH*-driven sfYFP and sfCFP are validated as robust tools for real-time symbiosis monitoring and for studying the effectiveness and competitiveness of rhizobia strains within the host, several caveats remain. Reporter performance is sensitive to physiological parameters such as oxygen, pH, and protein folding environment, with these constraints most notably affecting RFP performance in pea. Rare co-infection events indicate potential complexity in interpreting nodule occupancy, particularly in assays involving multiple strains. While automated imaging significantly improves throughput and reproducibility, ongoing benchmarking against standard approaches is necessary to ensure data robustness and accuracy. This effort is further constrained by the limited reference literature on the use of deep learning for quantifying nodulation traits.

While the systems reporter here shows clear efficacy for investigating rhizobium-legume symbiosis with fluorescent bioreporters, future optimization of fluorescent proteins, especially for red and far-red channels, will be essential to fully realize the potential of multiplexed visualization in diverse legumes. Integrating these improved reporters with conventional assessments and next-generation sequencing may provide a deeper understanding of plant-microbe interactions and symbiotic regulation. The combination of reliable promoter-reporter systems with scalable, automated quantification establishes a strong foundation for high-resolution, mechanistic studies of symbiotic nitrogen fixation and host-microbe ecology.

## Author contributions

BAG conceived the idea, and CKG, GS, ABP and MAS performed the experiments and analyzed the data. BAG, BMP and AHR supervised the work. All authors contributed to the writing of the manuscript.

## Acknowledgements

Funding for this work was provided by a New Innovator in Food & Agricultural Research (FFAR) grant to BAG ID: FF-NIA21-0000000061. BMP, BAG and ABP were funded by Specialty Crop Block Grant 22-242 from USDA/NIFA. Additional support for BMP was provided by hatch funds 7009493 and ND02438 from USDA/NIFA. BAG was further funded by hatch funds ND02439 from USDA/NIFA. The USDA-ARS partially supported this work under agreement No. 58-6064-3-011 (project No. FAR0037049). We acknowledge the technical support from research specialist Megan Ramsett, as well as Scott Hoselton from the Thomas Glass Biotech Innovation Core Lab at North Dakota State University.

## Supplementary Figures

**Supplementary Figure 1.**
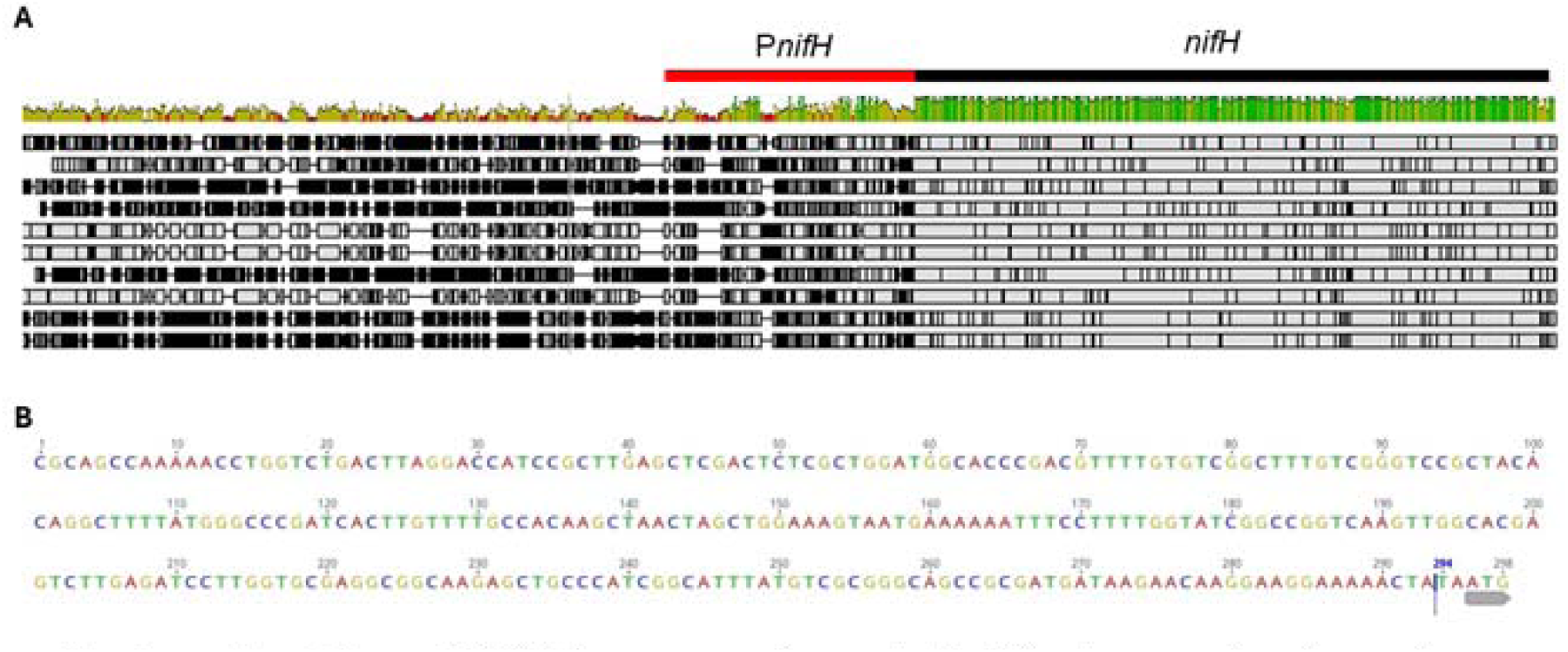
Multiple sequence alignment of *nifH* upstream regions from reference strains from several symbiotic *Mesorhizobium* species. These included *M. amorpha* CCNWGS0123. *M ciceri* bv *bisemdae* WSM1284 and WSM1271, *M jarvisii* ATCC33669, *M. cicen* CC1192A: *M muleiense* AHPC3O, *M. loti* TONO, and *M. loti* MAFF303099.

**Supplementary Figure 2.**
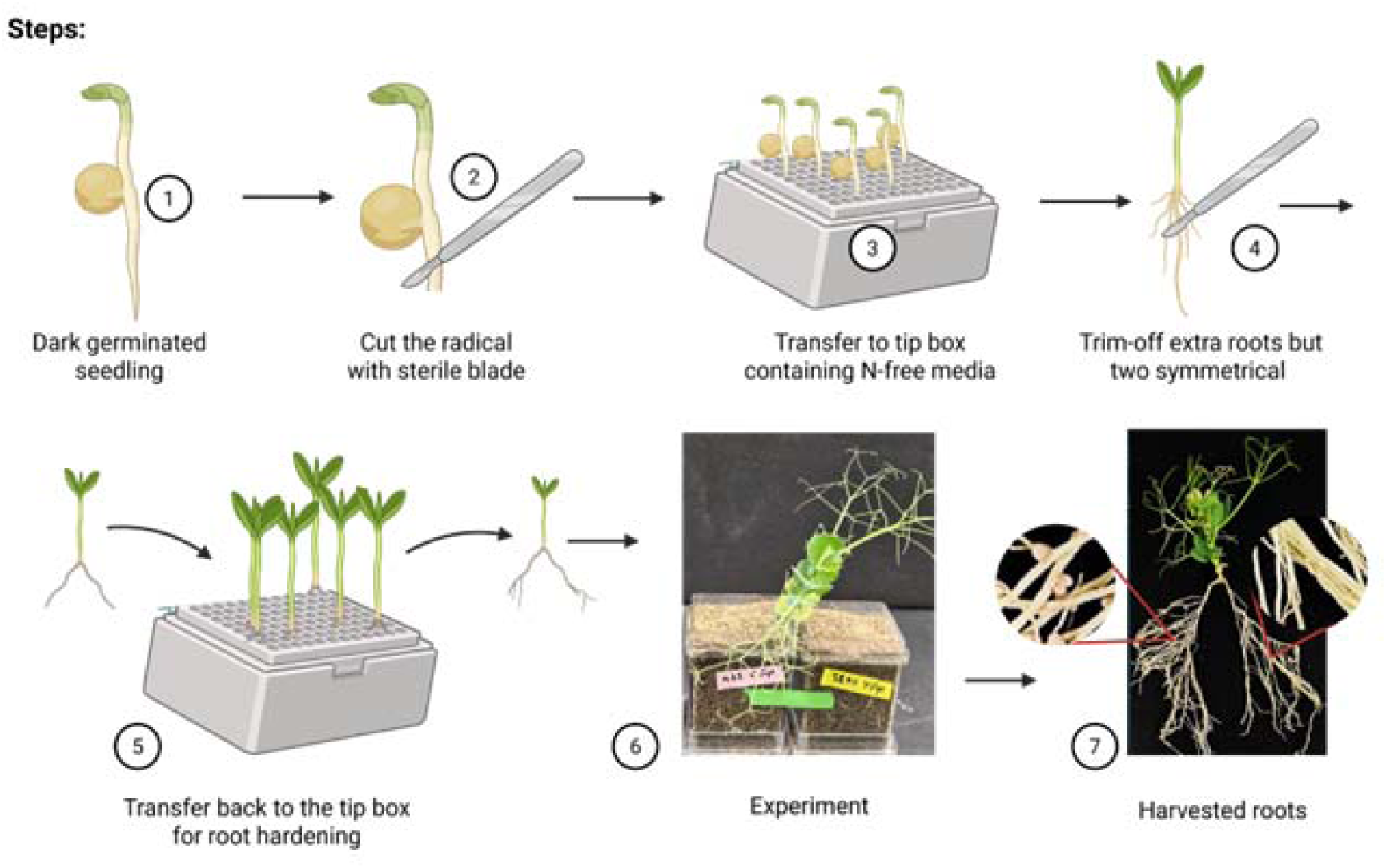
Split-root assay workflow. Flow diagram illustrating the split-root assay workflow used to study plant-microbial interactions. Two uniquely fluorophore-labeled *Rhi=obium* strains were inoculated separately into Leonard jars, each containing one-half of the split root system.

**Supplementary Figure 3.**
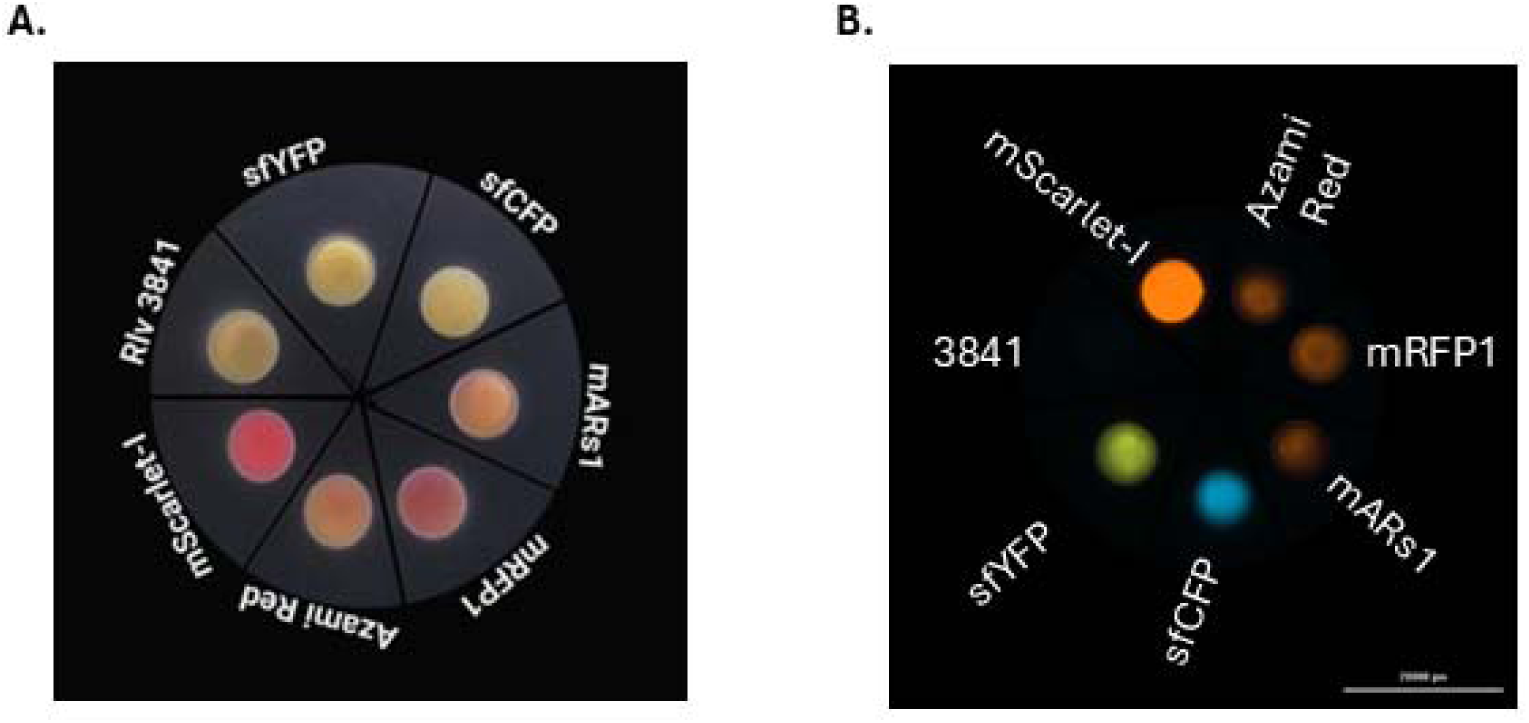
P*uni*-driven fluorescent labeling of *R. leguminosamm* on TY agar plates. (A) White-light image of ***R.*** *leguminosarum* strains labeled with different P11ni-driven fluorophores, including unlabeled Rlv3841 as the negative control. (B) *CytationTMj* imaging of *R. legwninosamm* strains (from Fig. Suppl. Fig. IA) labeled with different Puni-driven fluorophores, including unlabeled Riv3841 as the negative control.

**Supplementary Figure 4.**
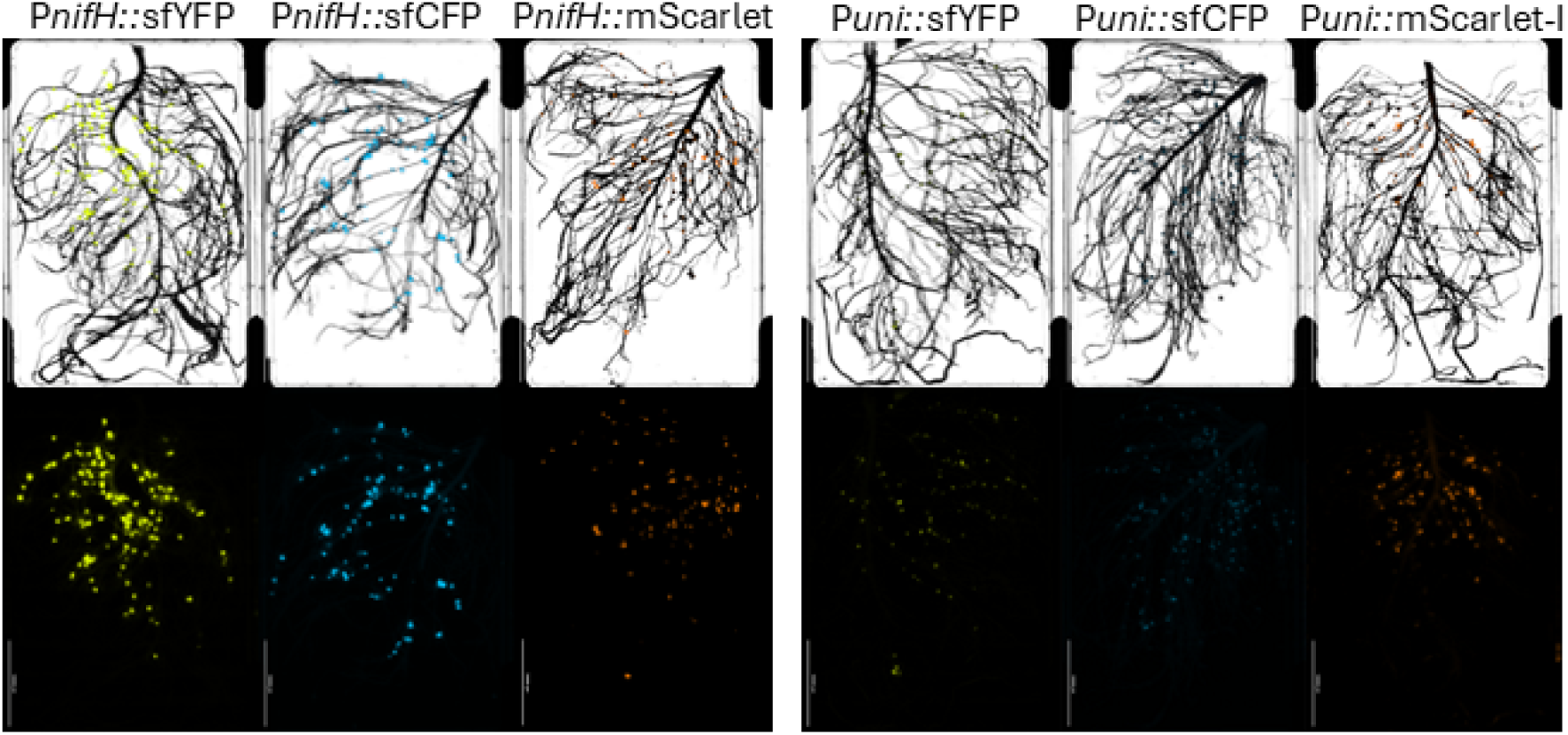
*Cytation^TM^ 5* imaging of *P. sativum* nodules inoculated with *R. l,egu11tiJ1osarum* strains expressing P*nifH-* and P*uni*-driveo fluorescent proteins (sfYFP, sfCFP, and mScarlet-1). Fluorescence images were captured using specific imaging parameters optimized for each fluorophore. Bright field (BF) and fluorescence settings were as follows: I. Excitation/emission wavelengths (nm): YFP (500/542), RFP (530/593), CFP (445/510) II. Light-emitting diode (LED) intensity: BF 10, YFP 10, RFP 10, CFP 10 III. Integration time (ms): BF 7, YFP 5, RFP 5, CFP 5 IV. Gain settings: BF 0.8, YFP 19.5, RFP I, CFP 24 A single set of parameters was maintained for each fluorescent protein across all samples.

**Supplementary Figure 5.**
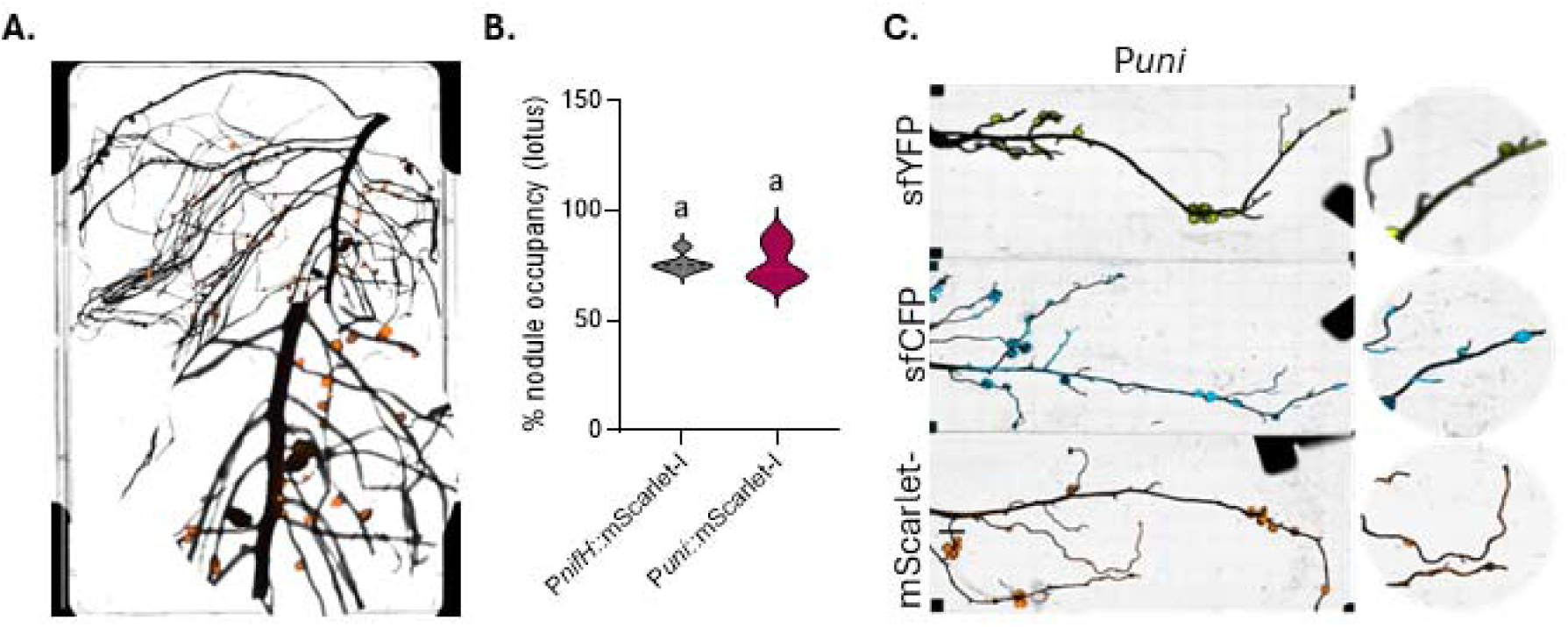
P*uni*-driveo expression of fluorescent proteins io *P. sativum* and *L. japonicus*. (A) *Cytation™ 5* imaging of *P. sativum* root nodules inoculated with *R. leguminosarum* expressing P*uni:*:mScarlet-I. (B) Percentage of fluorescent nodules exhibiting mScarlet-I fluorescence driven by either P*nifH* or *Puni* promoters in *P. sativum.* Data represent fiveindependent biological replicates. (C) *Cytation™ 5* imaging of *L.japonicus* root nodules inoculated with *Mjaponicum* labeled with Puni-driven sfYFP, sfCFP, and mScarlet-1. Distinct letters above bars indicate statistically significant differences (P < 0.05) as determined by one-way ANOVAfollowed by Tukey’s HSD test.

**Supplementary Figure 6.**
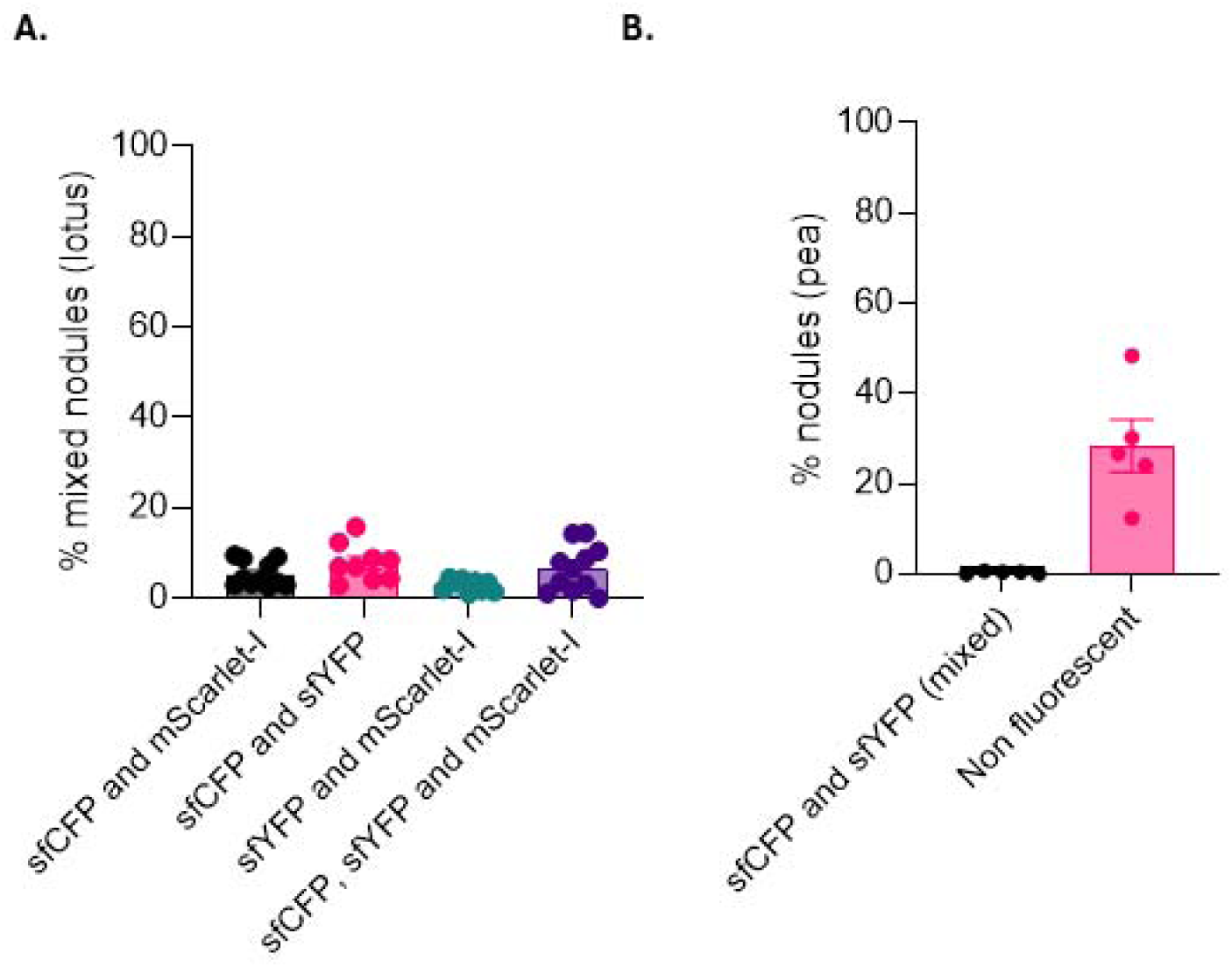
Percentages of root nodules containing mixed fluorescent labels in *P. sati,,mn* and *L. japo11icus* following c.o-inocnlation with rhizobial strains exressing distinct fluorescent tags. (A) Proportion of mixed nodules in *L. japonicus* roots co-inoculated with *Mjaponicum* strains carrying equal ratios of the following: i. P*nifH*::sfCFP and P*nifH*::mScarlet-1 ii. P*nifH*::sfCFP and P*nifH*::sfYFP iii. P*nifH*::sfYFP and P*nifH*::mScarlet-1 iv. Three-strain mixture of P*nifH*::sfCFP, P*nifH::sfYFP,* and P*nifH*::mScarlet-1. (B) Proportion of mixed and non-fluorescent nodules in *P. sativum* roots co-inoculated with *R. leguminosamm* strains expressing P*nijH*::sfCFP and P*nijH::sfYFP* at equal ratios. Data represent six to nine independent biological replicates for Fig. S6A, and five for Fig. S6B. Mixed nodules were identified by the simultaneous presence of multiple distinct fluorescent signals.

**Supplementary Figure 7.**
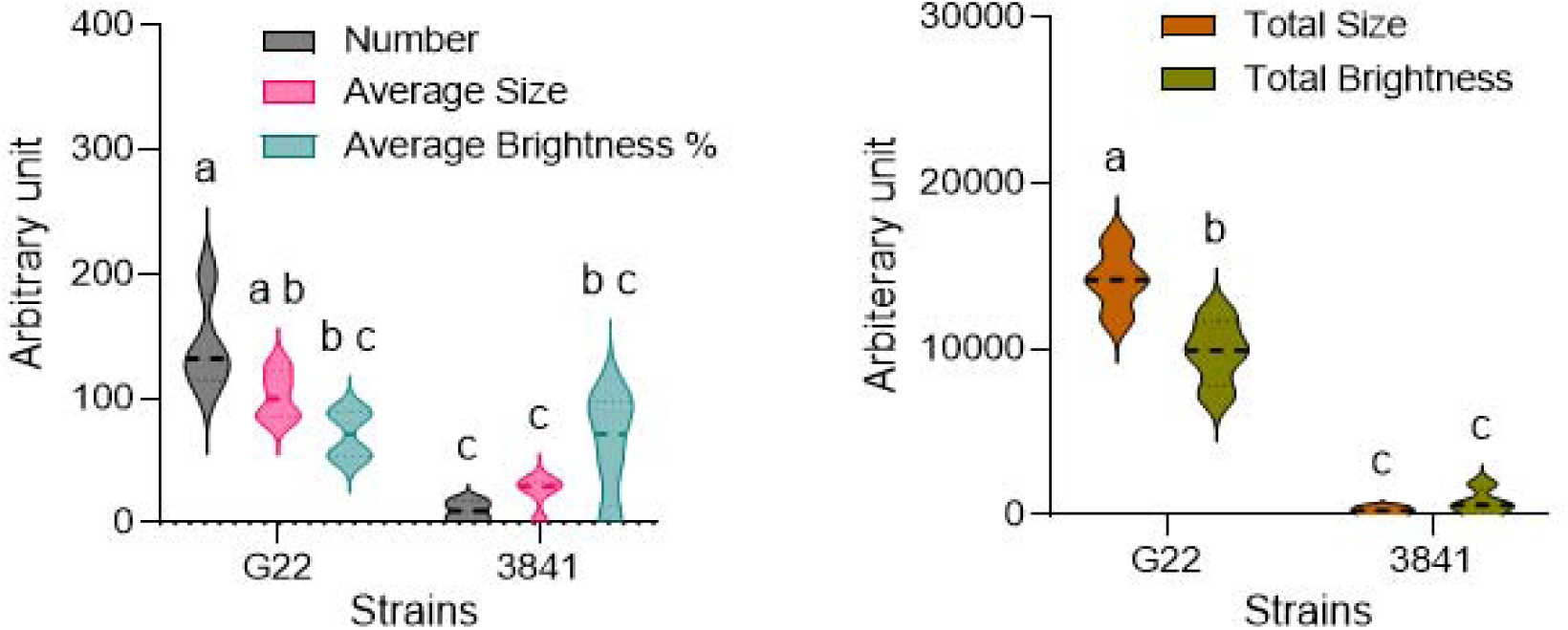
ImageJ-based quantification of nodulation traits from the split-root experiment shown in Fig. 5. Data were analyzed from four independent biological replicates. Nodulation traits such as nodule number, size, and brightness were quantified using ImageJ’s particle analysis tools. Data represent four independent biological replicates. Distinct letters above bars indicate statistically significant differences (P < 0.05) determined by two-way ANOVA followed by Tukey’s HSD test.

## Supplementary Tables

**Table S1:**
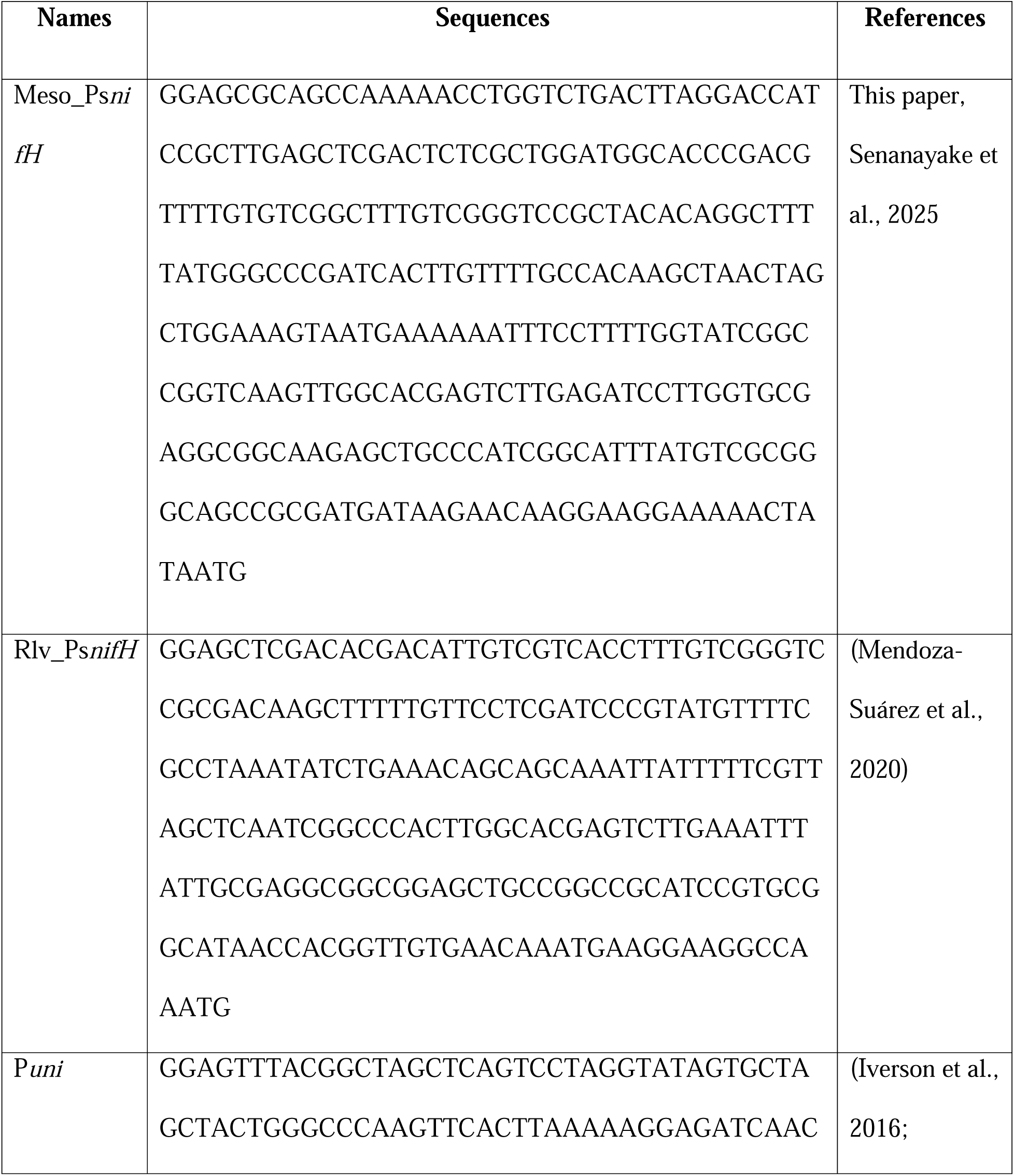
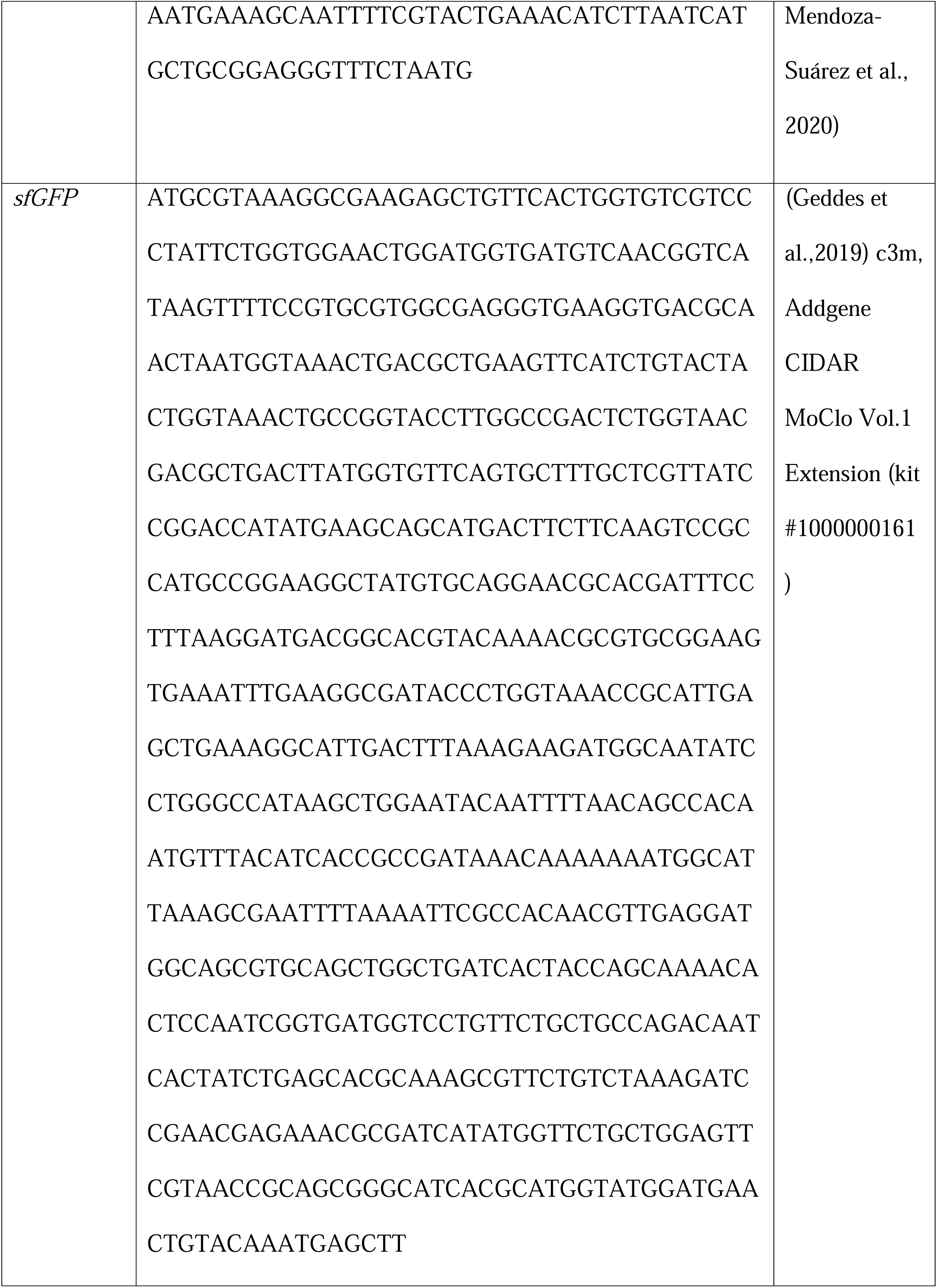
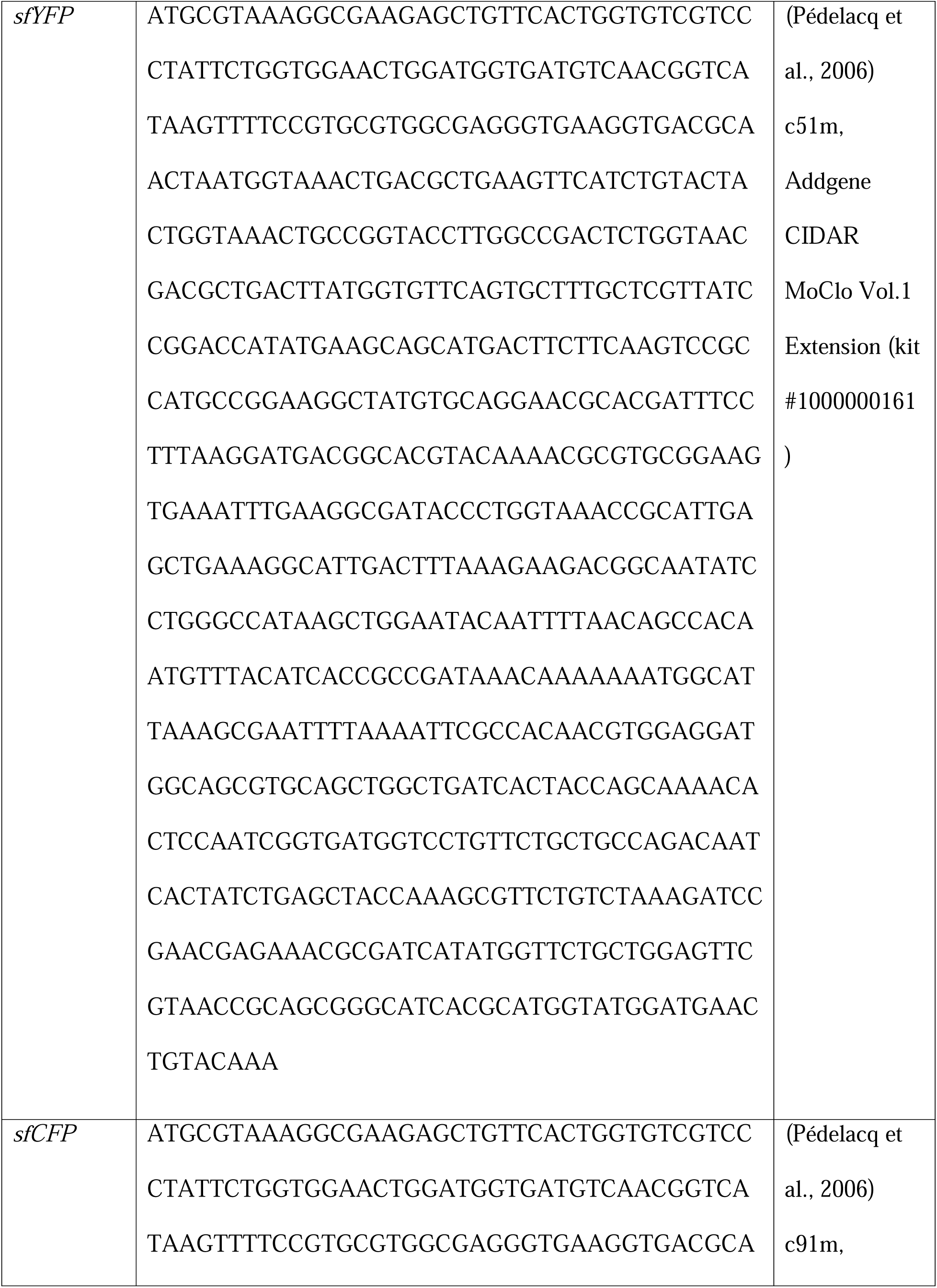
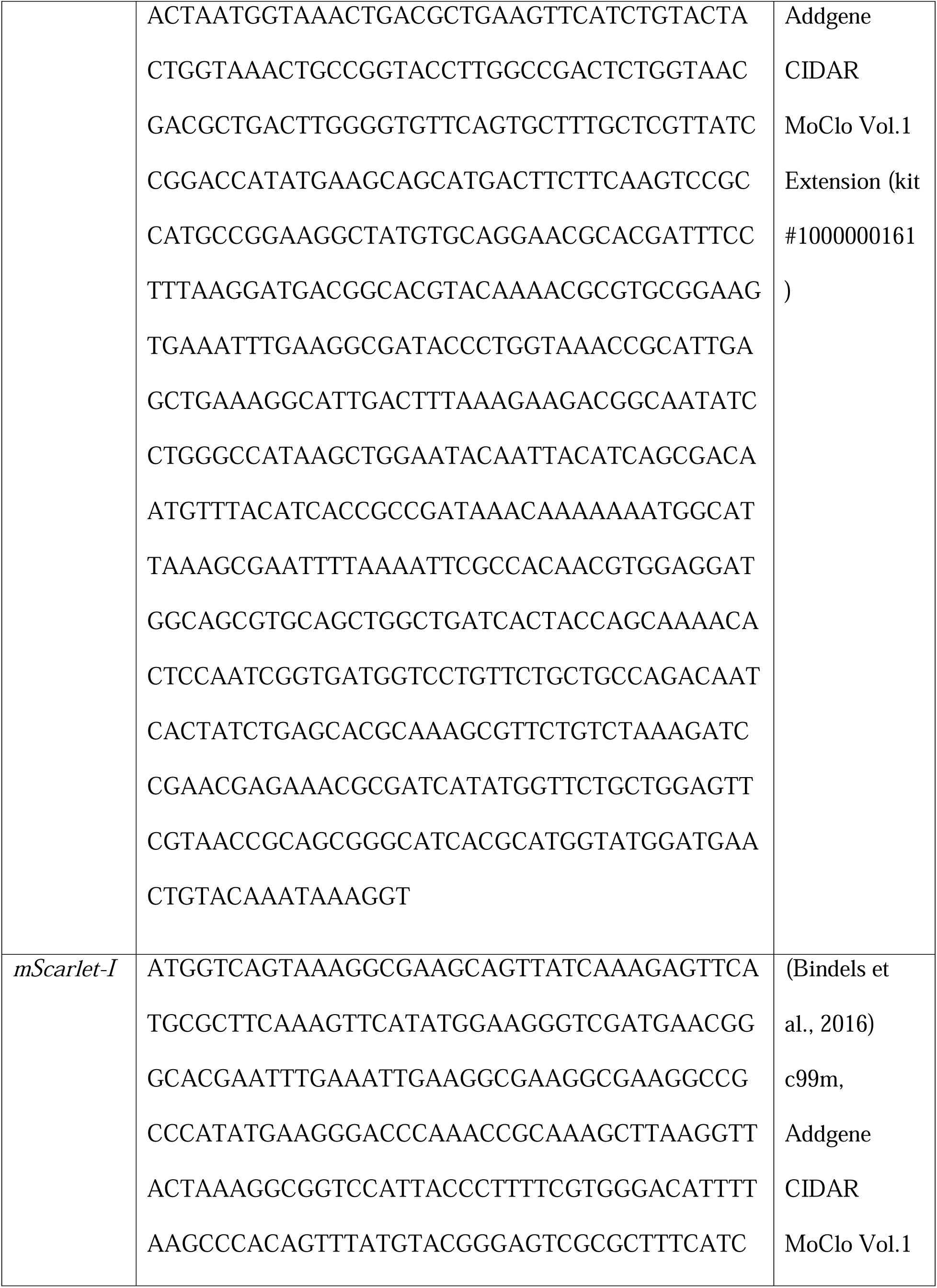
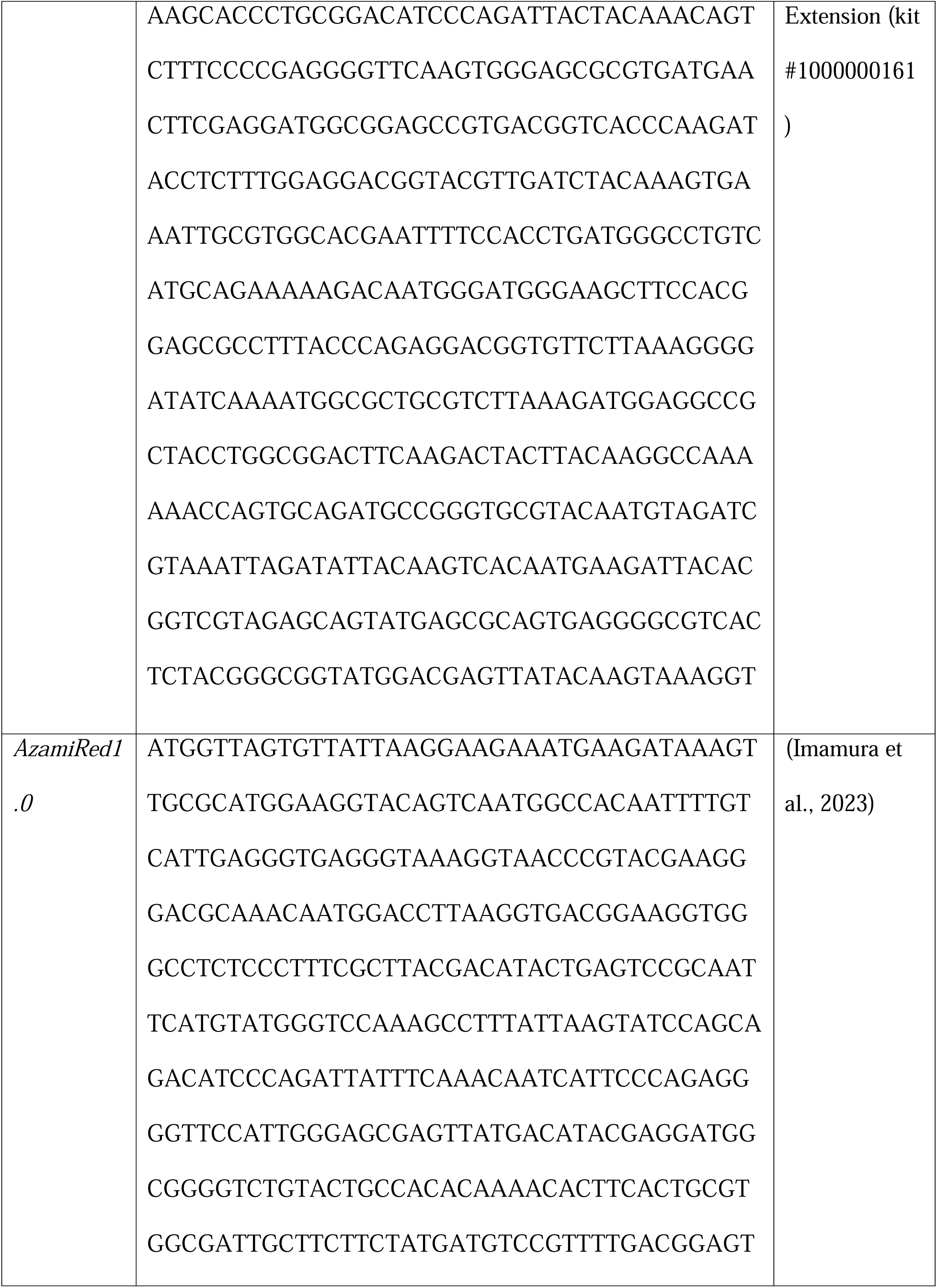
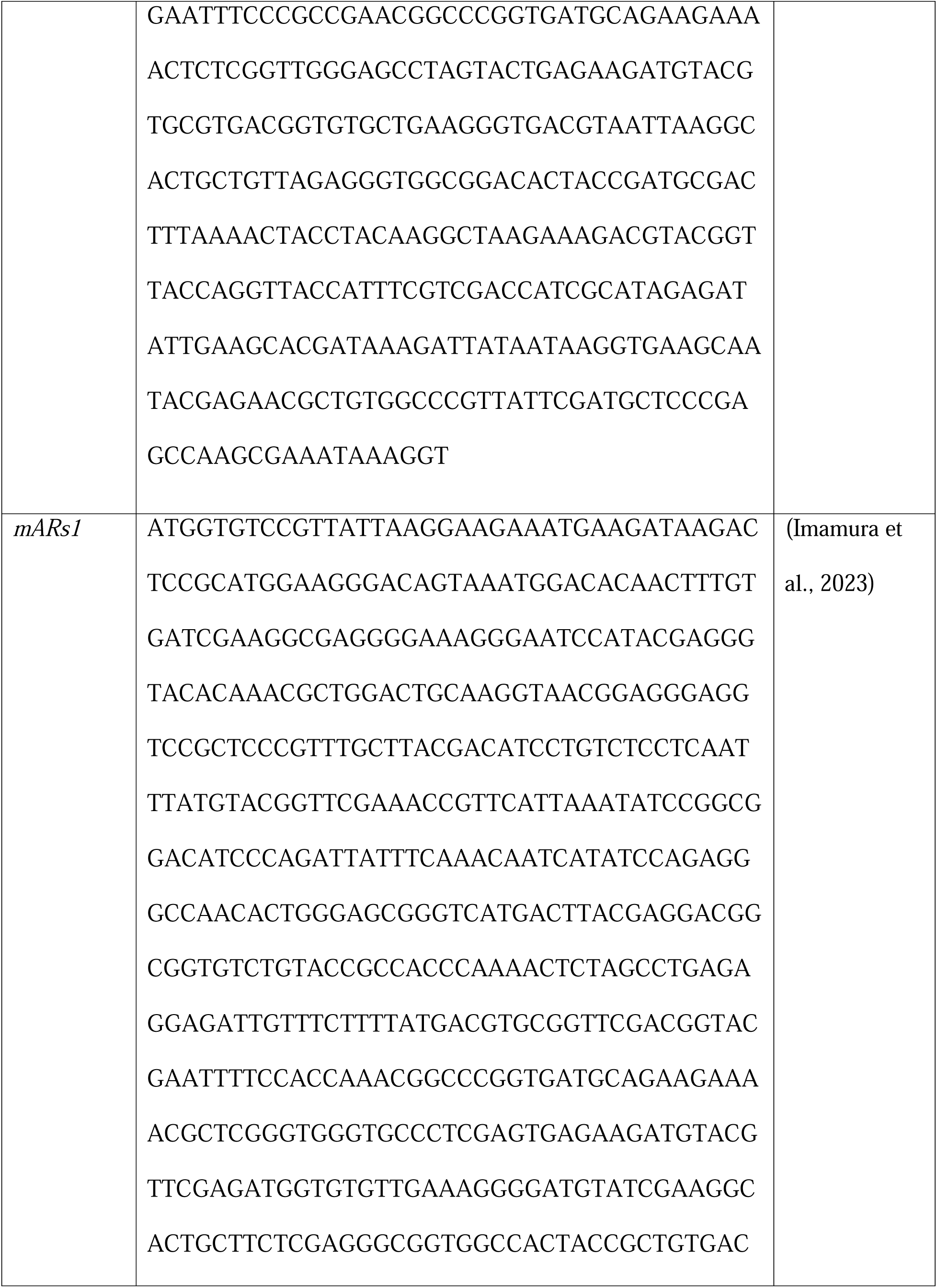
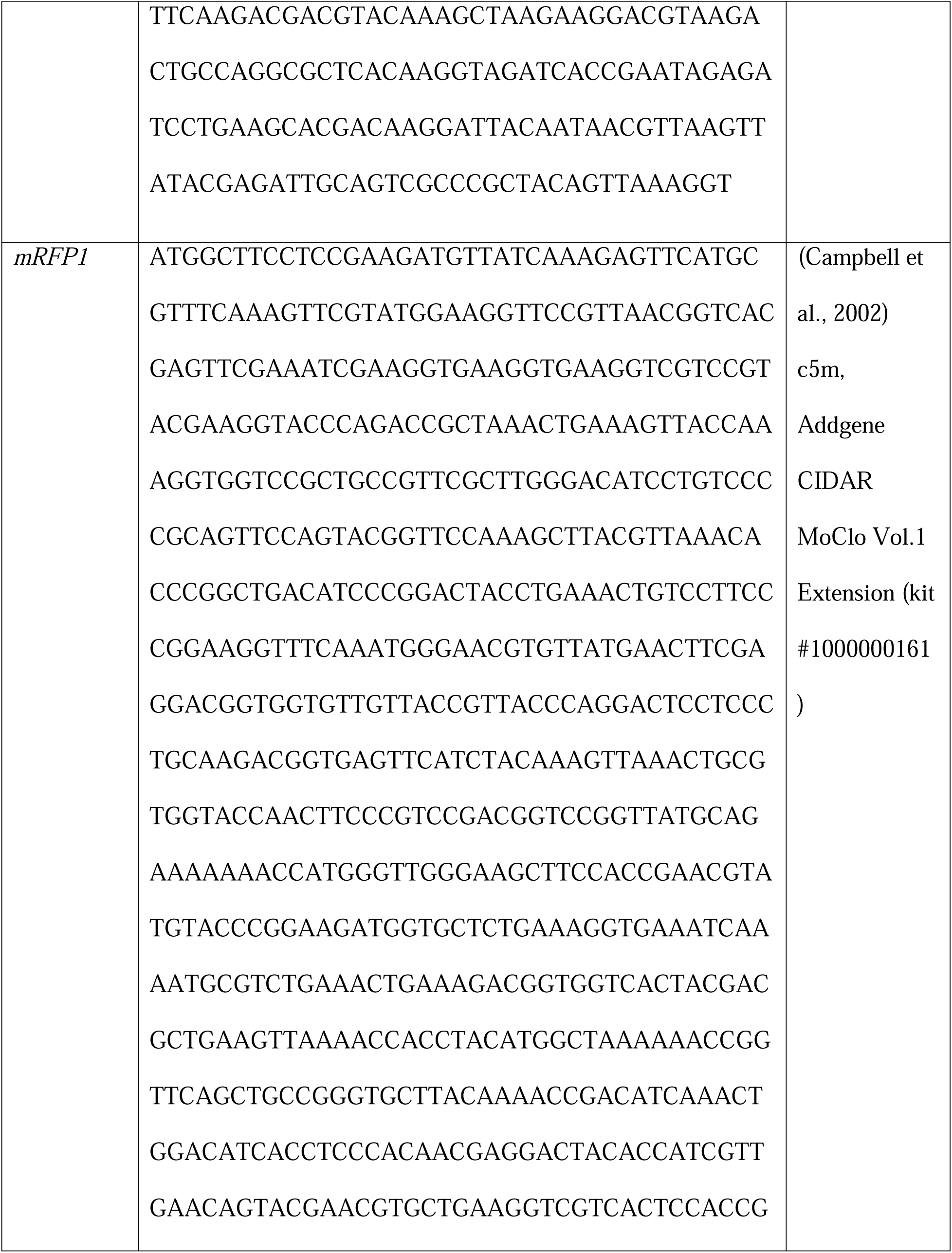
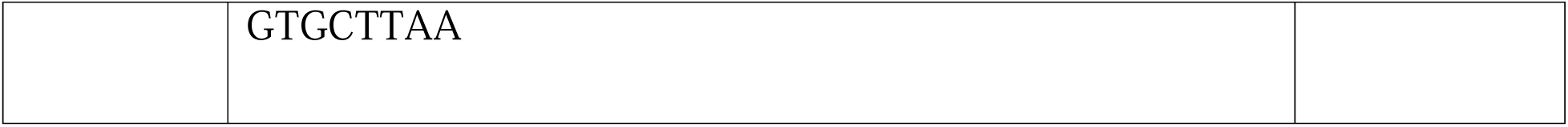
Nucleotide sequence of promoters and fluorescent protein encoding genes.

**Table S2.** The nitrogen-free rooting solution was prepared as previously described (47) with the following composition:

| Component | Concentration |
| --- | --- |
| $\text{CaCl}_2 \cdot 2\text{H}_2\text{O}$ | 1 mM |
| KCl | 100 $\mu\text{M}$ |
| $\text{MgSO}_4 \cdot 7\text{H}_2\text{O}$ | 800 $\mu\text{M}$ |
| Fe-EDTA | 10 $\mu\text{M}$ |
| $\text{H}_3\text{BO}_3$ | 35 $\mu\text{M}$ |
| $\text{MnCl}_2 \cdot 4\text{H}_2\text{O}$ | 9 $\mu\text{M}$ |
| $\text{ZnCl}_2$ | 0.8 $\mu\text{M}$ |
| $\text{Na}_2\text{MoO}_4 \cdot 2\text{H}_2\text{O}$ | 0.5 $\mu\text{M}$ |
| $\text{CuSO}_4 \cdot 5\text{H}_2\text{O}$ | 0.3 $\mu\text{M}$ |
| $\text{KH}_2\text{PO}_4$ | 50 g/L |
| $\text{Na}_2\text{HPO}_4$ | 56.8 g/L |

The pH of the solution was adjusted to $7.0 \pm 0.2$ using NaOH or HCl before use.

**Table S3.** The Jensen’s media was prepared as previously described (61) with the following composition:

| **Jesnsen's Media + 0.5 mM nitrogen (10x for 1 L)** | **Weight** |
| --- | --- |
| Calcium phosphate dibasic (CaHPO<sub>4</sub>) | 10 g |
| Potassium phosphate dibasic (K<sub>2</sub>HPO<sub>4</sub>) | 2 g |
| Magnesium sulfate heptahydrate (MgSO<sub>4</sub>· 7H<sub>2</sub>O) | 2 g |
| Ferrous chloride anhydrous (FeCl<sub>2</sub>) | 1 g |
| Potassium Nitrate (KNO<sub>3</sub>) | 505 mg |
| 1000x trace mineral solution | 10 mL |
| **Trace Mineral Solution (1000x per 1 L)** | **Weight** |
| Biotin | 0.4 g |
| CuSO<sub>4</sub> | 0.5 |
| H<sub>3</sub>BO<sub>3</sub> | 1 g |
| MnCl<sub>2</sub>· 4H<sub>2</sub>O | 0.5 g |
| Na<sub>2</sub>EDTA | 10 g |
| Na<sub>2</sub>MoO<sub>4</sub> | 1 g |

The pH of the solution was adjusted to $7.0 \pm 0.2$ using NaOH or HCl before use.

**Table S4.**
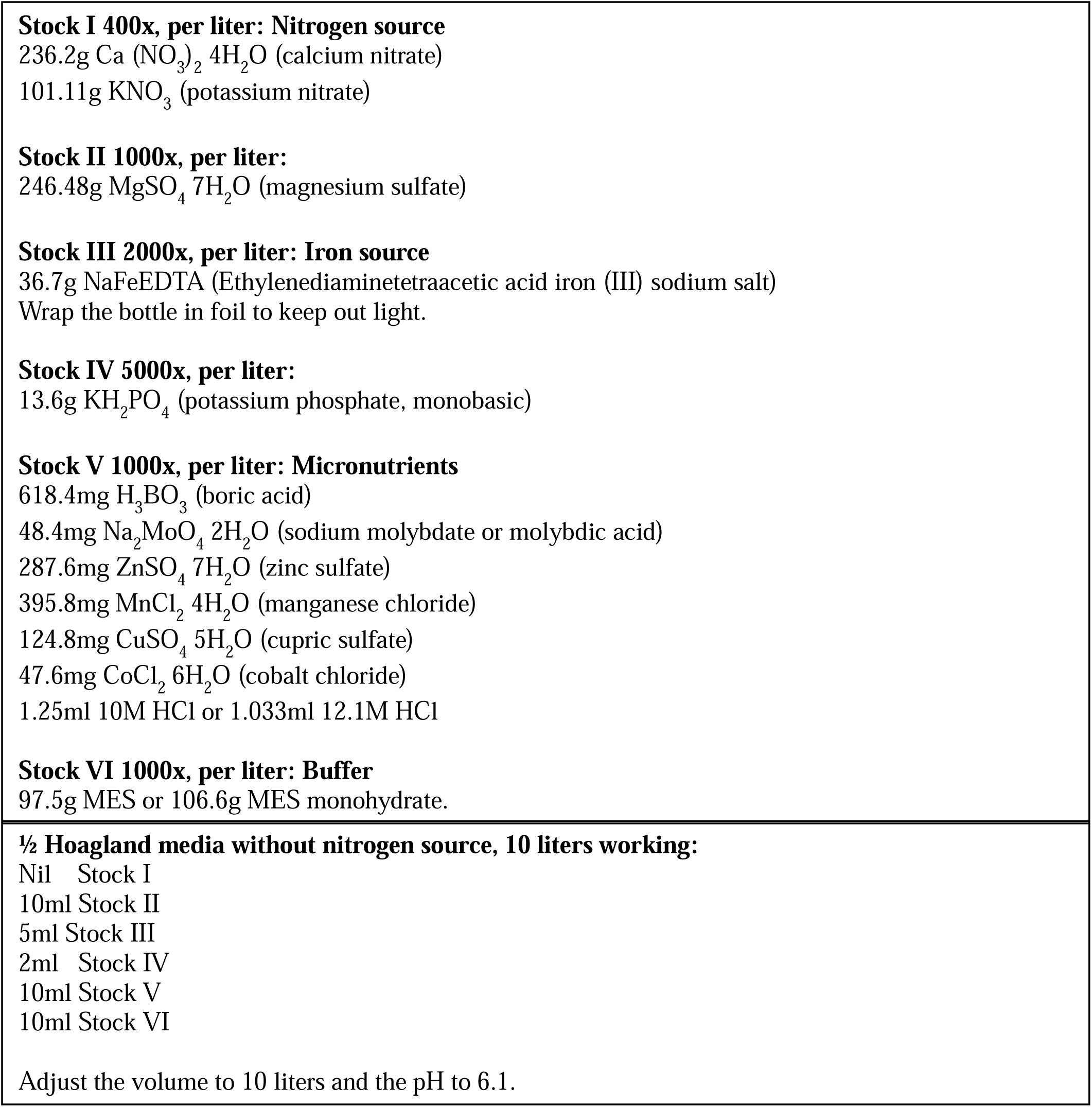
Composition of ½ Hoagland’s media, modified from Hoagland and Arnon 1950 (46):

